# DNA-damage dependent isoform switching modulates RIF1 DNA repair complex assembly and phase separation

**DOI:** 10.1101/2024.10.29.619708

**Authors:** Adenine Si-Hui Koo, Weiyan Jia, Sanghwa Kim, Mark Scalf, Claire E. Boos, Yuhong Chen, Demin Wang, Andrew F. Voter, Aditya Bajaj, Lloyd M. Smith, James L. Keck, Christopher J. Bakkenist, Lin Guo, Randal S. Tibbetts

**Author notes:** These authors contributed equally (Adenine Si-Hui Koo, Weiyan Jia).

## Abstract

How RIF1 (RAP1 interacting factor) fulfills its diverse roles in DNA double-strand break (DSB) repair, DNA replication, and nuclear organization remains elusive. Here we show that alternative splicing (AS) of a cassette exon (Ex32) encoding a Ser/Lys-rich (S/K) cassette in the RIF1 C-terminal domain (CTD) gives rise to RIF1-Long (RIF1-L) and RIF1-Short (RIF1-S) isoforms with different functional characteristics. We demonstrate that RIF1-Ex32 splice-in is mediated by an exonic splicing enhancer that is recognized by the splicing factor SRSF1 and antagonized by splicing inhibitors SRSF3 and SRSF7. Exposure to DNA damage inhibited Ex32 splice-in, potentiated the association of SRSF3 and SRSF7 with RIF1 pre-mRNA, and caused an increase in RIF1-S protein expression, which was also observed across a diverse set of primary cancers. Isoform-specific proteomic analyses revealed RIF1-L preferentially associated with mediator of DNA damage checkpoint 1 (MDC1) and sustained MDC1 focus formation to a greater extent than RIF1-S. We further show that the S/K cassette stabilized a novel phase separation activity of the RIF1 CTD and enhanced RIF1-L chromatin retention, which was reversed by CDK1-dependent phosphorylation of the RIF1 CTD in response to G_2_ DNA damage checkpoint inhibition. These combined findings suggest DNA damage-dependent RIF1 AS contributes to RIF1 functional diversification in genome protection.

## INTRODUCTION

Originally identified as a telomere-binding factor in *Saccharomyces cerevisiae*, the RIF1 (RAP1 interacting factor) gene encodes a ∼270 kDa protein that fulfills diverse roles in eukaryotic genome maintenance (1, 2). In mammals, RIF1 functions downstream of the canonical H2A.X-MDC1- 53BP1 signaling axis to influence DNA double-strand break repair (DSBR) pathway choice (3). In G_1_ and early S phase, 53BP1-dependent recruitment of RIF1 to DNA double-strand breaks (DSBs) promotes their repair via non-homologous end joining (NHEJ) while suppressing BRCA1- dependent homology-directed repair (HDR) (4–7). Specifically, RIF1 recruits Shieldin complex (SHLD1-3 and REV7) that suppresses 5’ → 3’ end resection to inhibit HDR (8–11). RIF1-deficient cells phenocopy elements of 53BP1 deficiency, including NHEJ and class-switch recombination (CSR) defects, as well as the inappropriate recruitment of BRCA1 to DSBs in G_1_ phase (4, 12–14). Furthermore, RIF1 deficiency rescues HDR defects and PARP inhibitor sensitivity of BRCA1- mutant cells, establishing RIF1 as a key player in 53BP1-dependent DSBR pathway choice (4, 12).

RIF1 also executes several important roles in DNA replication control. RIF1 is a component of the Bloom’s (BLM) helicase complex that suppresses deleterious end resection at stalled replication forks (RFs) to facilitate RF restart and recovery (15). RIF1 also delays replication origin firing through the recruitment of protein phosphatase 1 (PP1) to late origins, leading to dephosphorylation of the MCM4 subunit of the replicative minichromosome maintenance (MCM) DNA helicase (16). Therefore, RIF1-deficient cells exhibit delayed DNA replication rates, hypersensitivity to DNA replication inhibitors, and a defect in the IR-induced intra-S phase checkpoint (17, 18).

Possibly independent of its proximal role in suppressing origin firing, RIF1 also regulates genome-wide DNA replication timing (RT) through its effects on nuclear architecture, in which it organizes chromatins into topologic domains with similar RT in early G_1_ phase prior to the assembly of functional origin recognition complexes (19). RIF1-deficient mammalian cells or yeasts exhibit spatial changes in DNA replication domains that correlate with premature replication origin firing (20–22) (reviewed in Ref.(23)). RIF1 chromatin occupancy correlates with the RT of individual chromatin domains suggesting a role for RIF1 in bundling co-regulated origins (24). Interestingly, RIF1 participation in spatial organization of replication domains is genetically separable from its participation in RT regulation (25). Lastly, RIF1 also plays a role in the resolution of ultrafine DNA bridges in anaphase cells (26, 27).

How RIF1 simultaneously mediates DSBR pathway choice, replication origin regulation, and nuclear architecture maintenance is unclear. However, it is likely that protein isoforms, modular protein-protein interactions, and pathway-specific posttranslational modifications (PTMs) contribute to RIF1 functional diversity. The N-terminus of RIF1 (∼1–935 aa) comprises an array of 21 folded HEAT (Huntingtin, elongation factor 3 (EF3), protein phosphatase 2A alpha subunit and yeast PI3K TOR1) repeats that exhibits phosphorylation-dependent binding to 53BP1 and binding to SHLD3 (11, 28). The partially-folded carboxyl-terminal domain (CTD) contains three evolutionarily conserved regions (CR1 – CR3) based on sequence identity amongst vertebrate RIF1 proteins (15). CR2 additionally bears homology to the CTD of bacterial RNA polymerase α subunits (αCTD) (15). The RIF1 CTD mediates oligomerization, DNA binding, PP1 recruitment (through RVSF motif in CR1), and association with the BLM complex (15, 28–32). However, the full extent of its contributions to various RIF1 functions is unknown.

Here we show that RIF1 undergoes DNA damage-dependent alternative splicing (AS) to produce RIF1-L and RIF1-S isoforms which differ with respect to a 26 aa S/K cassette in the RIF1 CTD.

RIF1 AS was regulated by a suite of RNA-binding proteins including SRSF1, SRSF3, and SRSF7 whose expression and occupancy on RIF1 pre-mRNA changed in response to DNA damage and in primary cancers. We show that the S/K cassette enhanced RIF1-dependent accumulation of MDC1 at sites of DNA damage; strengthened RIF1 binding to chromatin; and promoted stable phase separation of the RIF1 CTD. By contrast, the DNA damage-induced RIF1-S isoform showed transient phase separation activity and weakly associated with chromatin, suggesting it is a more mobile RIF1 isoform. These studies illuminate mechanisms and functional consequences of RIF1 AS in response to DNA damage signaling to execute its roles in maintaining genome integrity.

## RESULTS

### RIF1 undergoes DNA damage and cell cycle dependent AS

The human RIF1 is a 2472 aa protein of approximately 270 kDa (Fig. 1A). RIF1 is expressed as two splice variants, RIF1-L and RIF1-S, that differ by the absence or presence of Exon 32 (Ex32). Ex32 encodes a 26 aa peptide which we have dubbed the S/K cassette owing to the predominance of Ser and Lys residues (Fig. 1A). RT-PCR analysis using primers flanking RIF1- Ex32 (Fig. 1B) revealed that RIF1-L (the 2472 aa isoform), was the major mRNA species in H- 460, U-2 OS, and HEK293T cells while HeLa cells had approximately equal amounts of both RIF1 isoforms (Fig. 1C). Cellular exposure to the radiomimetic drug Calicheamicin γ1 (CLM) increased relative abundance of the RIF1-S isoform in all four cell lines suggesting that DNA double-strand breaks (DSBs) repress Ex32 inclusion and promote the formation of RIF1-S (Fig. 1C). The decreased abundance of RIF1-L mRNA over the same time suggests this isoform may be selectively degraded under conditions of DNA damage. By contrast, the AS of a cassette exon in the Ubiquitin E3 ligase TRIP12 was not inhibited following CLM exposure in U-2 OS cells (Fig. 1D). Ex32 exclusion and repressed RIF1-L transcript production was CLM dose dependent and reached a maximum at 4-6 hours after treatment in HeLa (Fig. 1E) and U-2 OS cells (Fig. 1F). In consistent with RIF1 transcript quantification, RIF1-L/S protein ratio started to decrease at 4 hours post-CLM treatment in HeLa cells (Fig. 1G,H). Ionizing radiation (IR) dose-dependently promoted Ex32 skipping in H460 cells, indicating that Ex32 exclusion is a general response to DNA damage (Fig. 1I,J). RIF1 AS also fluctuated during the cell cycle, with the highest RIF1-L/RIF1-S mRNA ratio in G_2_/M-phase and the lowest ratio in G_1_-phase HeLa cells (Fig. 1K,L), suggesting that RIF1 AS is regulated by cell cycle signaling.

**Figure 1.**
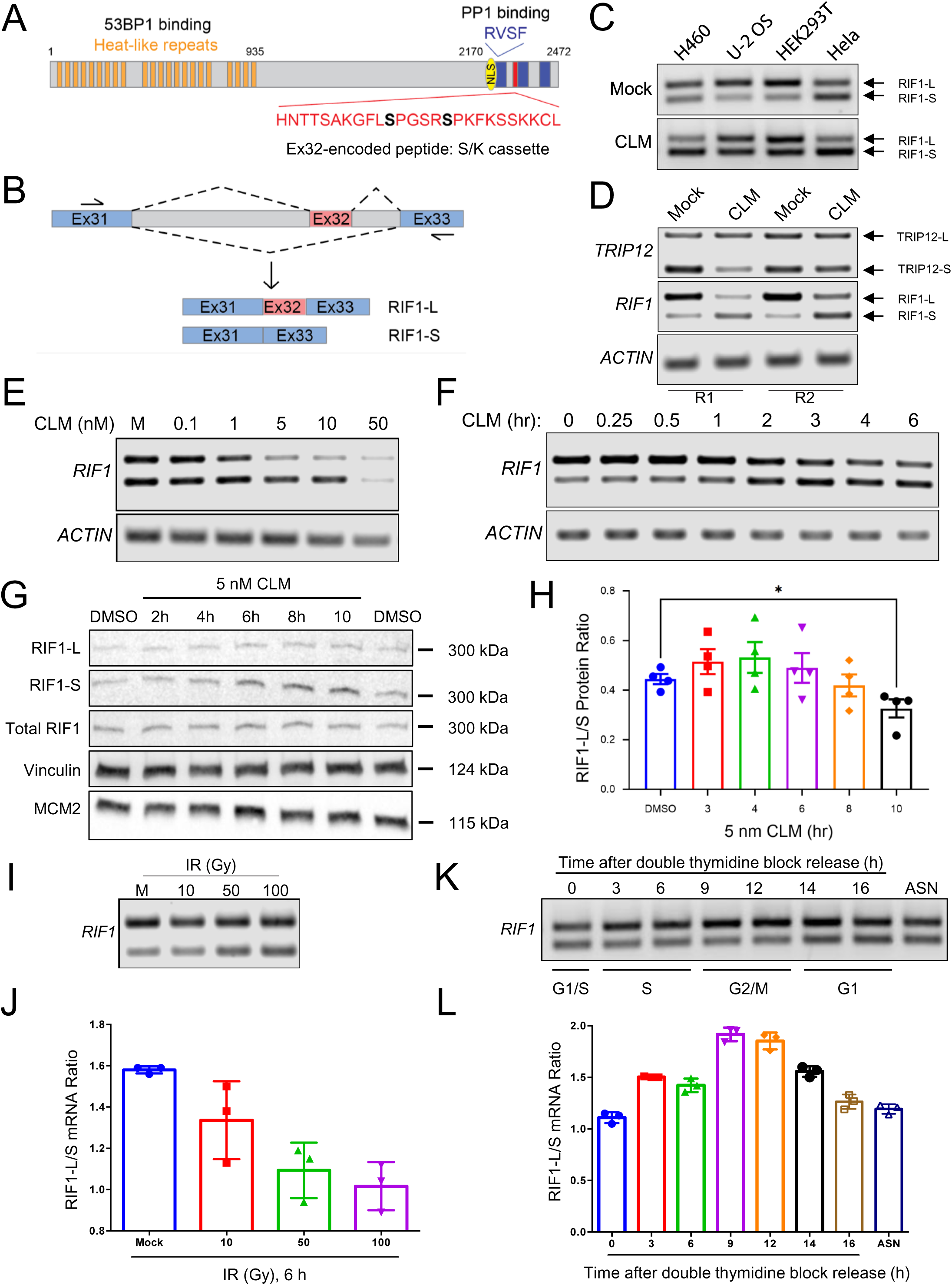
RIF1 undergoes DNA damage and cell cycle dependent AS. (A) RIF1 protein domains. Blue rectangles from left to right correspond to CR1, CR2 and CR3 respectively. Ex32- encoded S/K cassette is highlighted in red with its sequence shown. CDK1 phosphorylation sites (S2260 and S2265) are in bold. (B) RIF1 Ex32 splicing assay with a forward primer targeting Exon 31 (Ex31) and a reverse primer targeting Exon 33 (Ex33). (C) Exposure to the radiomimetic drug Calicheamicin γ1 (CLM) (10 nM for 6 h) reduced the RIF1-L/S mRNA ratio in H460, U-2 OS, HEK293T, and HeLa cells. (D) Differential response of RIF1 and TRIP12 AS to CLM. U-2 OS cells were treated with 10 nM CLM for 6 h and the total mRNA was analyzed for Ex32 and Ex3 inclusion in RIF1 and TRIP12, respectively. (E) CLM dose dependence of RIF1 splicing regulation in HeLa cells treated with the indicated CLM concentration for 4 h. (F) Time dependence of RIF1 splicing regulation by 10 nM CLM in U-2 OS cells. (G, H) RIF1-L/S protein ratio in HeLa cells treated with 5 nM CLM for the indicated timepoints. The mean RIF1-L/S protein ratio ± standard error was calculated by densitometry. Each dot represents an individual biological replicate, N = 4 (*. p ≤ 0.05; from repeated measures one-way ANOVA with Geisser-Greenhouse correction and Dunnett’s multiple comparisons test). (I, J) Dose dependence of ionizing radiation (IR)- induced repression of Ex32 inclusion in H460 cells 6 h after exposure. The mean RIF1-L/S mRNA ratio ± standard error was calculated by densitometry. Each dot represents an individual biological replicate, N = 3. (K, L) RIF1 AS in HeLa cells fluctuates during the cell cycle. HeLa cells were released from a double-thymine block and harvested at the indicated times. The mean RIF1-L/S mRNA ratio ± standard error was calculated by densitometry. Each dot represents an individual biological replicate, N = 3.

We tested the impacts of canonical DNA repair inhibitors on CLM-dependent Ex32 skipping. While RIF1-L/RIF1-S ratios were comparable between CLM-treated cells with vehicle, PARP, or ATM inhibitors, the combination of CLM and a DNA-PK inhibitor potentiated Ex32 skipping, which may be indicative of enhanced DSB induction (Sup. Fig. 1A,B). These combined findings indicate that RIF1-Ex32 splicing is dynamically regulated by genotoxic stress and during the cell cycle.

### RIF1 isoform usage is altered in cancer

We queried publicly available RNA-Seq data from The Cancer Genome Atlas (TCGA) to estimate the abundance of RIF1-L and RIF1-S transcripts in normal versus tumor tissue across four selected cancer types by IsoformSwitchAnalyzeR (33) (Fig. 2A). Total RIF1 mRNA expression was significantly downregulated in breast invasive carcinoma (BRCA) relative to the matched normal breast tissue, while RIF1 expression levels were relatively upregulated in colon adenocarcinoma (COAD), lung adenocarcinoma (LUAD), and lung squamous carcinoma (LUSC) (Fig. 2B). When focusing on the isoform levels, RIF1-S mRNA expression was significantly upregulated in BRCA, COAD, LUAD, and LUSC while RIF1-L expression was only significantly downregulated in BRCA but remained unchanged in COAD, LUAD, and LUSC (Fig. 2C). Strikingly, the usage of RIF1-S and RIF1-L isoforms in all four cancer subtypes showed the same trends: RIF1-S isoform usage was significantly increased, while RIF1-L isoform usage was significantly decreased, except in COAD, where RIF1-L isoform usage was slightly reduced, though not statistically significant (Fig. 2D). These findings suggest that an RIF1 isoform switch from RIF1-L to RIF1-S may be associated with primary cancers.

**Figure 2.**
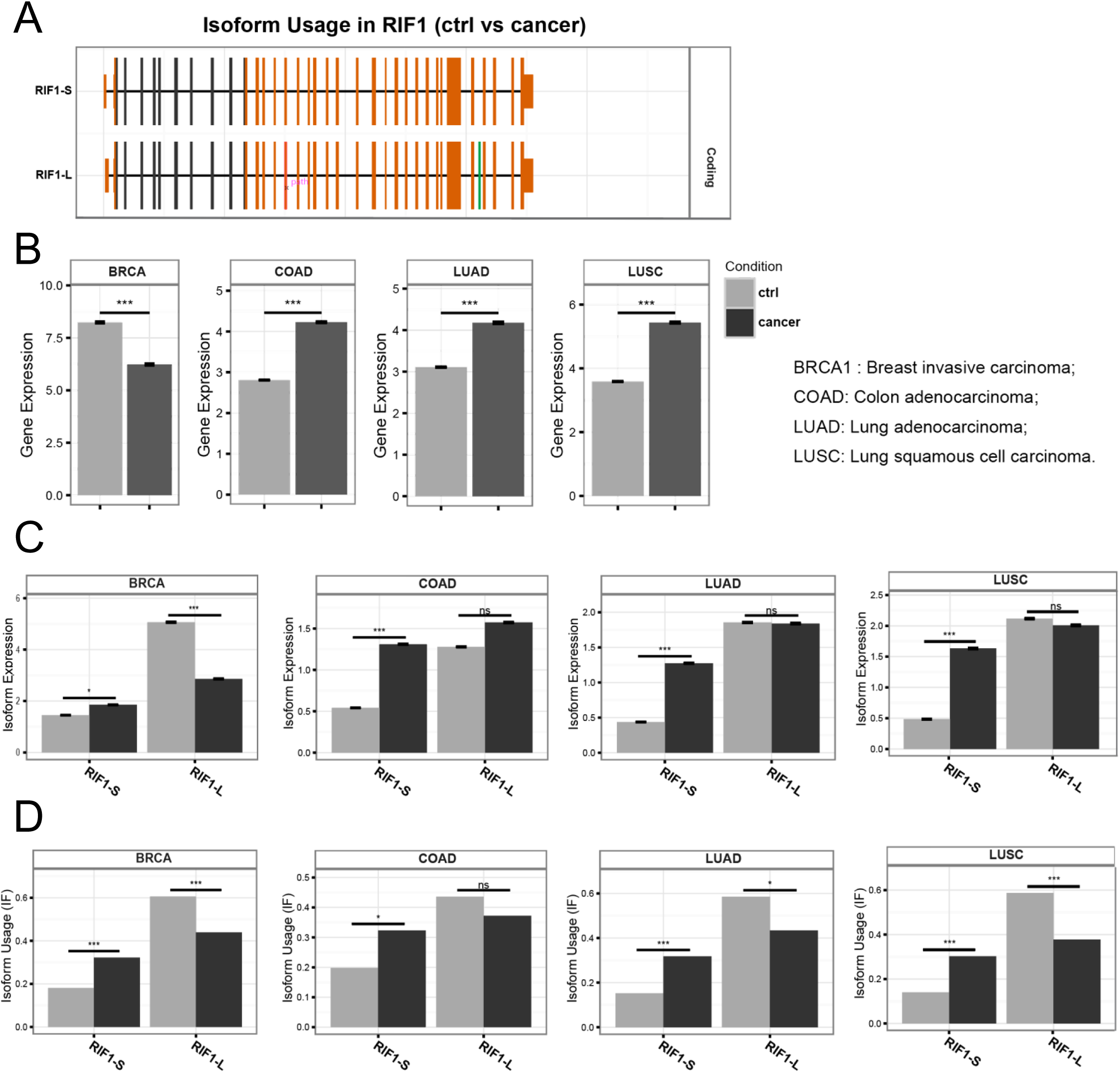
**RIF1 Ex32 splicing changes across cancer types**. (A) Exonic structure of RIF1-S and RIF1-L. (B-C) RIF1 isoform expression data was estimated from The Cancer Genome Atlas (TCGA) data through IsoformSwitchAnalyzeR (B) Total RIF1 gene expression in breast invasive carcinoma (BRCA) (Control, N = 114; Cancer, N = 1097), colon adenocarcinoma (COAD) (Control, N = 41; Cancer, N = 460), lung adenocarcinoma (LUAD) (Control, N = 59; Cancer, N = 516), and lung squamous cell carcinoma (LUSC) (Control, N = 51; Cancer, N = 502) (***. FDR < 0.001 from EdgeR differential expression analysis and two-tailed Mann-Whitney test). (C) The proportion of RIF1-S transcript relative to RIF1-L transcript in all four cancer types (ns. not significant; *. FDR < 0.05; ***. FDR < 0.001 from EdgeR differential expression analysis and two-tailed Mann-Whitney test). (D) RIF1 isoform usage (ns. not significant; *. FDR < 0.05; ***. FDR < 0.001 from EdgeR differential expression analysis and two-tailed Mann-Whitney test).

### Point mutations in Ex32 disrupt its inclusion into RIF1 transcripts

We used CRISPR/Cas9 to disrupt Exon 2 (*RIF1^-/-^*), or Exon 32 (*RIF1-L^-/-^)* in U-2 OS cells. We generated two *RIF1^-/-^*lines (H1 and 2C5); two *RIF1-L^-/-^* lines (A6 and 2A2) containing frameshift mutations in Ex32 leading to the creation of a premature stop codon and likely encoded for truncated RIF1-L proteins; and a third *RIF1-L^-/-^* line (H11) harboring a homozygous 15 nt deletion that removes amino acids 2261-2265 (Fig. 3A). Interestingly, all three *RIF1-L^-/-^* cell lines showed dramatic reductions in RIF1-L transcript and a corresponding increase in RIF1-S transcript while mutations in Ex2 (*RIF1^-/-^*) had no impact on the relative proportion of RIF1-L and RIF1-S transcript (Fig. 3B,C). As expected, RIF1-L protein was undetectable in *RIF1-L^-/-^* (2A2) U-2 OS cells while total RIF1 levels were only slightly reduced relative to controls (Fig. 3D,E). On the other hand, *RIF1-L^Δ5^* (H11) harboring an in-frame deletion showed a decrease in both RIF1-L and total RIF1 expression in relative to wild-type U-2 OS cells (Fig. 3D,E).

**Figure 3.**
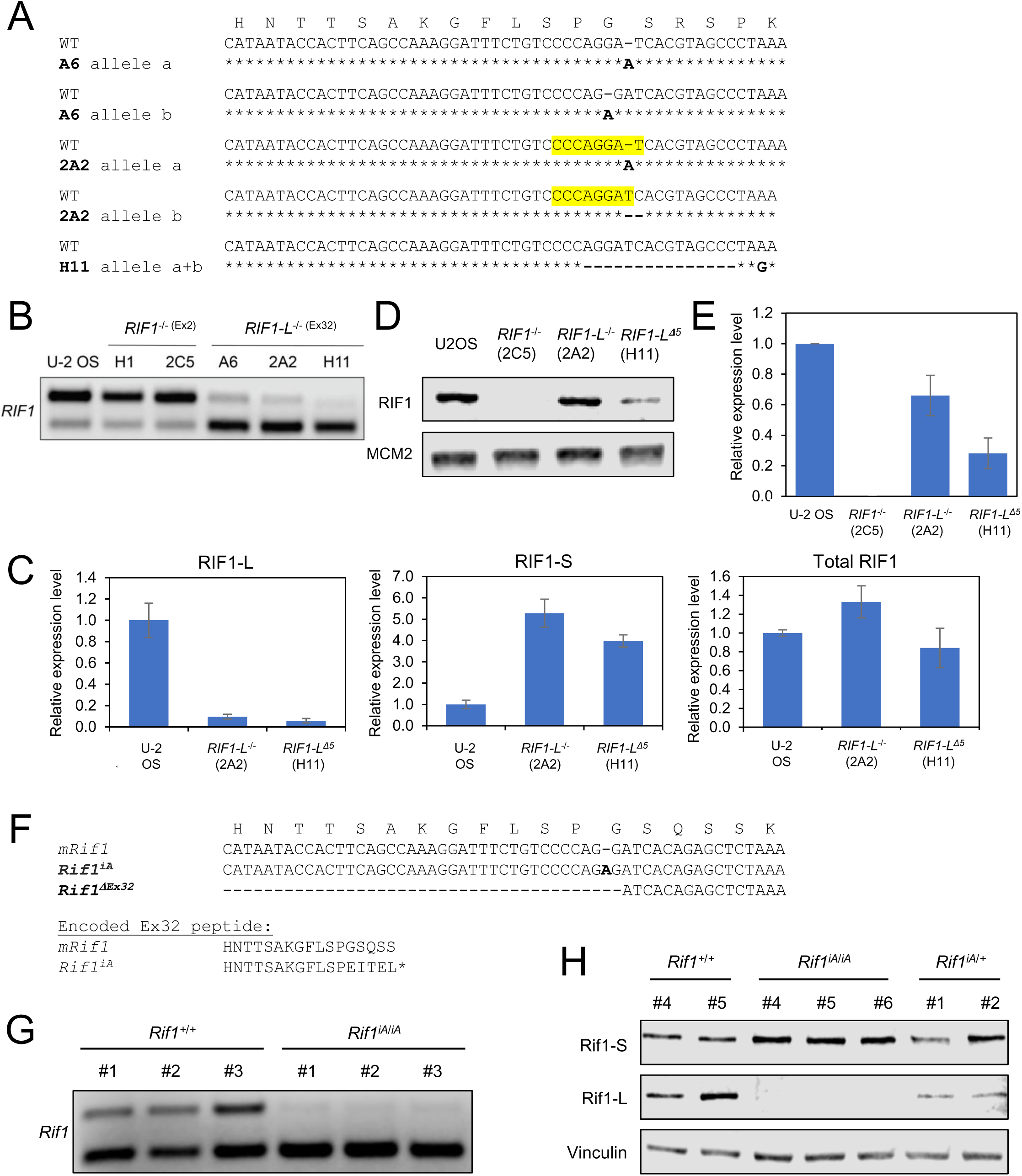
Point mutations in Ex32 diminish RIF1-L transcript formation. (A) Three *RIF1-L^-/-^*U-2 OS clones (A6, 2A2, and H11) harboring CRISPR-induced mutations (in bold) aligned to the wild-type (WT) RIF1-Ex32 allele. A6 and 2A2 have two distinct alleles “a” and “b” whereas H11 is homozygous. Disrupted SRSF1 binding site in Clone 2A2 was highlighted in yellow. (B, C) RT- qPCR showing reduced expression of RIF1-L mRNA and a corresponding increase in RIF1-S mRNA in *RIF1-L^-/-^* U-2 OS cell lines. Total RIF1 transcripts stayed relatively constant in WT and mutated U-2 OS cell lines. CRISPR-generated mutations in RIF1-Exon 2 (Ex2) (Clones H1 and 2C5), did not change RIF1-Ex32 AS. (D, E) Western blot analysis showed that *RIF1-L^-/-^*(Clone 2A2) and *RIF1-L^Δ5^* (Clone H11) have reduced total RIF1 expression while Clone 2C5 (*RIF1^-/-^*) has an undetectable RIF1 expression. (F) Murine *Rif1* alleles generated through CRISPR-mediated gene editing of Ex32. Rif1^iA^ mice harbor a single A insertion whereas Rif1^ΔEx32^ has a 129 nt deletion spanning Intron 31 and Ex32. (G) RIF1 splicing assay from three Rif1^+/+^ and Rif1^iA/iA^ mice showed reduced expression of RIF1-L and increased expression of RIF1-S mRNA in testis from homozygous Rif1^iA^ mice. (H) Western blot analysis of RIF1-L and RIF1-S protein expression in *Rif1^+/+^*, *Rif1^iA/+^*, and *Rif1^iA/iA^* testis extracts using isoform-specific antibodies showed a similar trend in (G).

### Point mutations in Ex32 disrupt its inclusion into RIF1 transcripts *in vivo*

We also generated two lines of mice selectively deficient for RIF1-L: a *Rif1^iA^* line harboring a single A insertion between G6604 and G6605 of the RIF1-L coding sequence and *Rif1^ΔEx32^*, which harbors a 129 nt deletion spanning the 3’ end of Intron 31 (In31) and the 5’ portion of Ex32 (T6477- G6605) (Fig. 3F). Similar to what was observed in U-2 OS cells harboring point mutations in Ex32, expression of RIF1-L mRNA was greatly reduced in testis extracts prepared from *Rif1^iA/iA^*mice, while RIF1-S mRNA was correspondingly upregulated, likely owing to a defect in Ex32 splicing (Fig. 3G). The fact that the iA mutation greatly reduced the RIF1-L/RIF1-S mRNA ratio supports the idea that Ex32 contains an exonic splicing enhancer (ESE) that is sensitive to small indels. Residual RIF1-L transcripts in *Rif1^iA/iA^* animals contain a premature termination codon in Ex32 and are likely substrates for the nonsense mediated mRNA decay (NMD) pathway. As expected, RIF1-L protein was undetectable in homozygous *Rif1^iA/iA^* mice while RIF1-S protein levels were upregulated relative to *Rif1^+/+^* mice (Fig. 3H).

Homozygous *Rif1^iA^* and *Rif1^ΔEx32^* mice were fertile, outwardly normal in appearance, and exhibited normal B and T cell development; normal proportions of mature B and T cells; and comparable rates of mitogen-induced B and T cells proliferation from two independent experiments, each with at least two mice per genotype (Sup. Fig. 2A-F). In contrast to what was observed in *Rif1*^-/-^ mice (4, 7), B cells from *Rif1^ΔEx32^* mice exhibited normal rates of class switching to IgG1, IgG2a, IgG2b, and IgG3 (Sup. Fig. 3A-D). Because RIF1-S accounts for all *Rif1* gene dosage in *Rif1^ΔEx32^* mice, these findings indicate that Ex32-encoded S/K cassette is not required for canonical roles of RIF1 in CSR.

### Identification of RIF1 splicing regulators

We carried out a siRNA screen of 151 siRNAs targeting RNA binding proteins (RBPs) for modulators of RIF1 AS in HeLa cells (Sup Fig. 4A-C) based on the prediction by RBPmap tool (https://doi.org/10.1093/nar/gku406). The primary siRNA screen and secondary shRNA validation screen implicated six RBPs whose silencing changed the RIF1-L/RIF1-S splicing ratio at least two-fold. Polypyrimidine tract-binding protein 1 (PTBP1), RNA Binding Motif Protein 28 (RBM28), serine and arginine-rich splicing factors (SRSFs) 3 and 7 were identified as the negative regulators of Ex32 inclusion, whereas small nuclear ribonucleoprotein U1 subunit 70 (snRNP70) and SRSF1 were identified as the positive regulators of Ex32 inclusion (Sup. Fig. 4C, Fig. 4A,B). The implication of SRSF1 as a RIF1 splicing factor is consistent with findings of Yu *et al.* who identified RIF1-Ex32 in a screen for SRSF1-regulated splicing events (34). We investigated SRSF2 as an additional putative regulator given that its binding site closely overlaps that of SRSF1 (35), and these two splicing factors often work in the same complex with snRNP70 (36). None of the identified RIF1-Ex32 splicing regulators significantly modulated RIF1-Ex1a splicing in the siRNA screen and shRNA knockdown (Fig. 4A).

**Figure 4.**
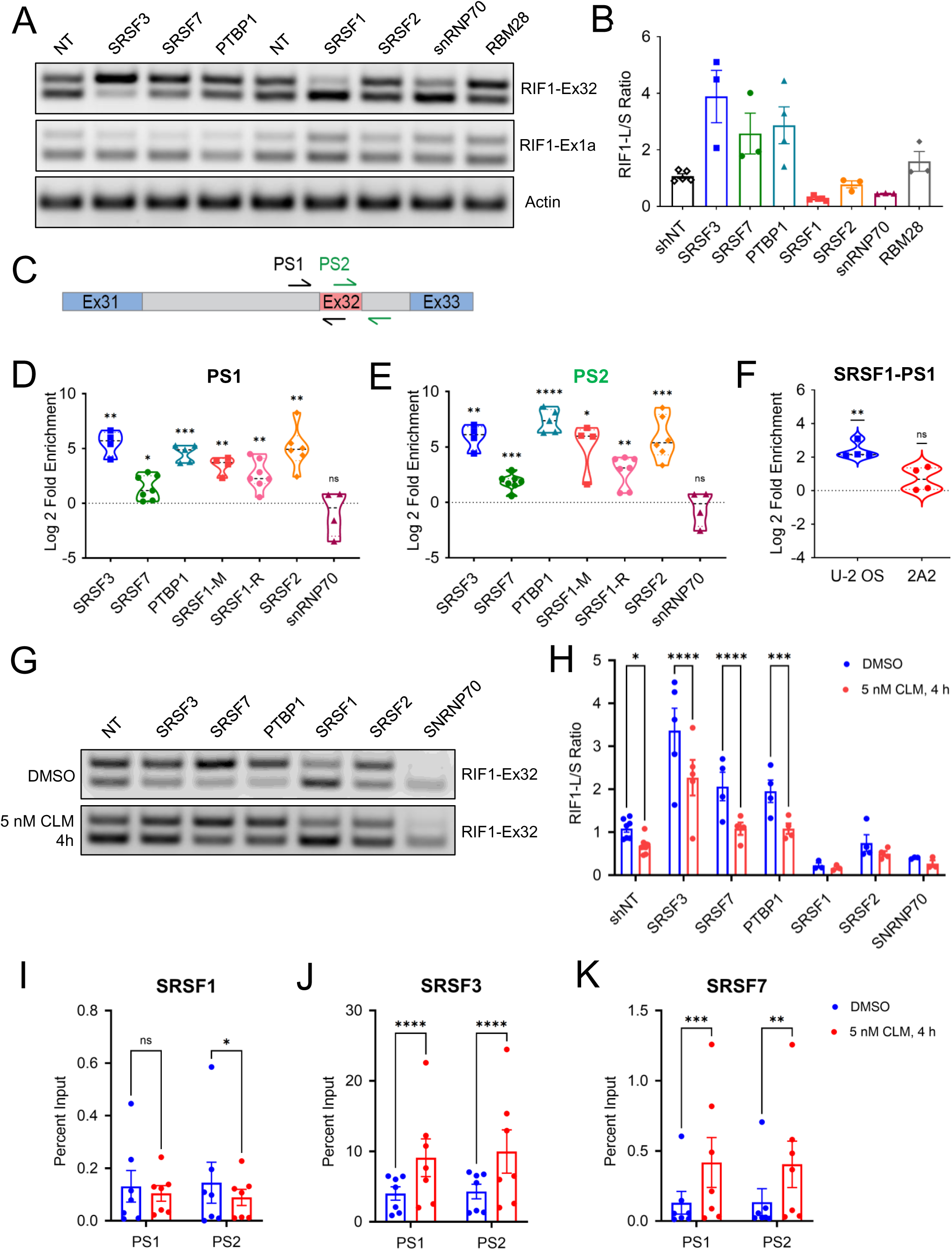
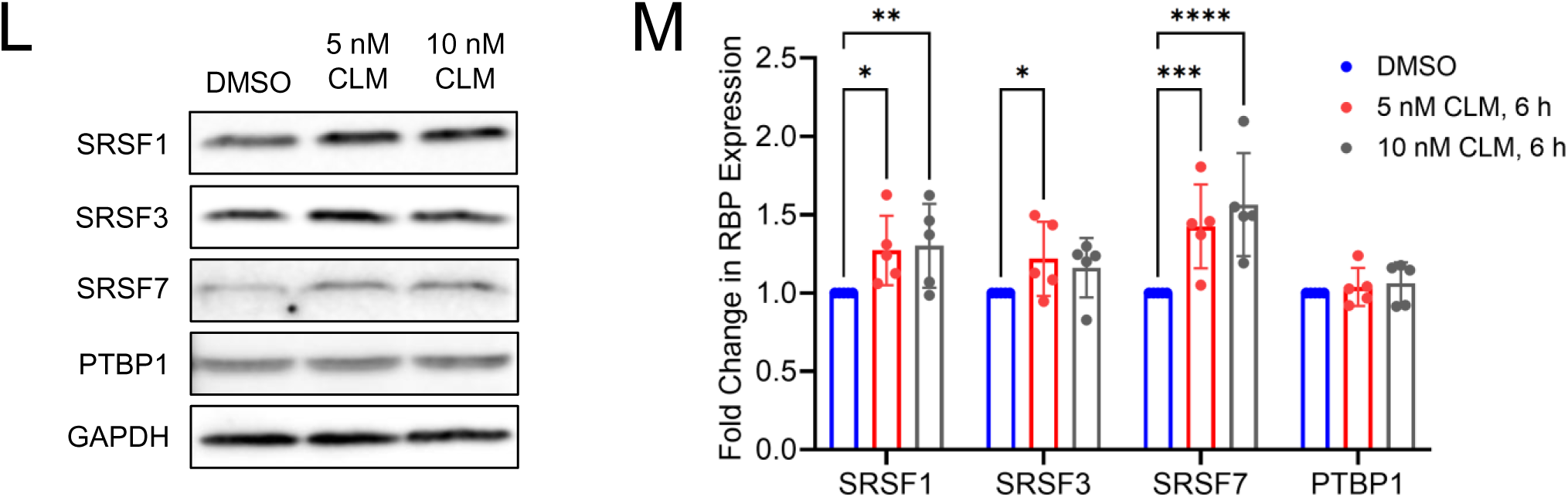
Identification of RBPs that regulate RIF1 AS in response to DNA damage. (A) Lentiviral shRNA vectors targeting putative RIF1 splicing regulators were transduced into HeLa cells to assess effects on RIF1-Ex32 and RIF1-Ex1a AS. Non-targeting (NT) shRNA served as a negative control. (B) Quantification of RIF1-L/S mRNA ratios from (A) based on densitometry. Each dot represents an individual biological replicate, bar height corresponds to mean ± standard error, 3 ≤ N ≤ 5. (C) Relative locations of the two intron-exon primer pairs used in RNA-IP qPCR experiments. PS1 targets RIF1-In31 and Ex32 whereas PS2 targets RIF1-Ex32 and In32. (D, E) Relative log2 fold enrichment of RIF1 pre-mRNA from native RNA-IP experiments in HeLa cells with antibodies targeting the indicated RBPs from PS1 and PS2 amplification as illustrated in (C). Median and interquartile range were shown as dotted lines, each dot represents an individual biological replicate, 4 ≤ N ≤ 7 (ns. not significant; *. p ≤ 0.05; **. p ≤ 0.01; ***. p ≤ 0.001; ****. p ≤ 0.0001 from two-tailed one sample t-test). (F) Relative log2 fold enrichment of RIF1 pre-mRNA from native RNA-IP experiments done in WT U-2 OS cells and Clone 2A2 (*RIF1-L^-/-^* U-2 OS, Fig. 3). Median and interquartile range were shown as dotted lines, each dot represents an individual biological replicate, N = 4 (ns. not significant; **. p ≤ 0.01 from two-tailed one sample t-test). (G) Representative gel image of HeLa cells transduced with the indicated lentiviral shRNA vectors and treated with 5 nM CLM for 4 h for RIF1-Ex32 splicing assay. (H) Quantification of RIF1-L/S mRNA ratios from (G) based on densitometry. Each dot represents an individual biological replicate, bar height corresponds to mean ± standard error, 3 ≤ N ≤ 5 (*. p ≤ 0.05; ****. p ≤ 0.0001 from repeated measures two-ways ANOVA and Šidák’s multiple comparisons test). (I-K) Percent input from native RNA-IP experiments with SRSF1, SRSF3, or SRSF7 in HeLa cells treated with DMSO or 5 nM CLM for 4 h. Each dot represents an individual biological replicate, bar height corresponds to mean ± standard error, N = 7 (ns. not significant; *. p ≤ 0.05; **. p ≤ 0.01; ***. p ≤ 0.001; ****. p ≤ 0.0001 from repeated measures two-ways ANOVA and Šidák’s multiple comparisons test). (L) Representative Western blot image of SRSF1, SRSF3, SRSF7, and PTBP1 expression level from DMSO, 5 nM and 10 nM CLM treatment for 6 h in HeLa cells. GAPDH was included as loading control. (M) Quantification of the fold change in normalized RBP expression from (L) relative to DMSO control. Each dot represents an individual biological replicate, bar height corresponds to mean fold change ± standard error, N = 5 (*. p ≤ 0.05; **. p ≤ 0.01; ***. p ≤ 0.001; ****. p ≤ 0.0001 from repeated measures two-ways ANOVA and Dunnett’s multiple comparisons test).

We next employed native RNA-IP to test whether candidate RIF1 splicing factors directly associated with RIF1 pre-mRNAs. HeLa cell extracts were immunoprecipitated with antibodies specific to SRSF3, SRSF7, PTBP1, SRSF1, SRSF2 and/or snRNP70 or the corresponding normal mouse and rabbit IgG controls. Following elution and purification, bound pre-mRNA was analyzed by RT-qPCR using two different intron-exon primer pairs, PS1 and PS2 as shown in Fig. 4C. While there was little to no enrichment of RIF1-Ex32 containing pre-mRNA from snRNP70- IPs, RNA-IP of all other splicing factors showed significant enrichment over IgG IP controls. In addition, both SRSF1 antibodies used (SRSF1-M and SRSF1-R) showed consistent and significant enrichment, strongly implicating SRSF1 as a direct Ex32 binding factor (Fig. 4D,E).

We noted that mutations that disrupted Ex32 inclusion into RIF1 mRNA transcripts in 2A2 and A6 U-2 OS cells were adjacent to a putative SRSF1 binding site (CCCAGGAT) as highlighted in Fig. 3A (35–39). Considering this, we compared SRSF1 binding to RIF1 pre-mRNA between *RIF1^+/+^* and *RIF1-L^-/-^* (Clone 2A2) U-2 OS cells. SRSF1 enrichment was not observed in 2A2 U-2 OS cells, suggesting that SRSF1 associates with Ex32 directly through the CCCAGGAT element (Fig. 4F).

### DNA damage induces the association of SRSF3 and SRSF7 with RIF1 pre-mRNA

We next sought to determine whether one or more negative regulators of Ex32 inclusion were targets of CLM-dependent regulation. We found that the knockdown of any single negative regulator was not sufficient to attenuate Ex32 skipping in CLM-treated Hela cells, suggesting potential redundant functions between these factors (Fig. 4G,H). On the other hand, native RNA-IP demonstrated a significant increase in SRSF3 and SRSF7 association with RIF1 pre-mRNA in CLM-treated HeLa cells (Fig. 4J,K). By contrast, SRSF1 association with RIF1 pre-mRNA slightly decreased following CLM treatment, though the effect was marginally significant (Fig. 4I). The inverse relationship between SRSF1 and SRSF3/SRSF7 binding to RIF1 transcripts plausibly explains Ex32 skipping in response to CLM. Based on these findings we evaluated whether CLM affected the localization (Sup. Fig. 5A) or the expression of SRSF1, SRSF3, SRSF7, and PTBP1 in response to CLM (Fig. 4L,M). While the nucleo-cytoplasmic localization of these RBPs did not show any obvious changes after CLM treatment (Sup. Fig. 5A), the total expression of SRSF1 and SRSF7 was significantly increased in CLM-treated HeLa cells (Fig. 4L,M). There was also a trend toward higher SRSF3 expression following CLM treatment; however, the difference was only marginally significant (Fig. 4L,M). PTBP1 expression was unaffected by CLM. Interestingly, a comparison of the median expression level of SRSF1, SRSF2, SRSF3, SRSF7, and PTBP1 in BRCA, COAD, LUAD, and LUSC subsets from the TCGA data revealed overexpression of SRSF3 and PTBP1, which may contribute to the low RIF1-L/RIF1-S ratio associated with these tumors (Fig. 2, Sup. Fig. 5B).

### RIF1 isoforms behave similarly in DNA replication control

To facilitate study of RIF1-L and RIF1-S isoforms, we reconstituted *RIF1^-/-^* U-2 OS cells with Dox-inducible, GFP-tagged, RIF1-L and RIF1-S cDNAs (see Materials and Methods) (Sup. Fig. 6A). Both *RIF1^-/-^*:GFP-RIF1-L and *RIF1^-/-^*:GFP-RIF1-S were targeted to IR-induced foci with qualitatively similar magnitude (Sup. Fig. 6B). GFP-RIF1-L and GFP-RIF1-S comparably suppressed MCM4 hyperphosphorylation, which is reflective of unscheduled replication origin firing (30) (Sup. Fig. 6C) and rescued the DNA replication patterning defect seen in *RIF1^-/-^* cells. Specifically, RIF1-L and RIF1-S restored the perinuclear and perinucleolar EdU incorporation patterns typical of mid-S phase cells that are almost completely absent in *RIF1^-/-^*cells (Sup. Fig. 6D-F) (40). Given the basic nature of the S/K cassette and its proximity to the RIF1 DNA binding domain, we measured DNA binding affinity of purified RIF1-L and RIF1-S C-terminal domains (RIF1^CTD^-L and RIF1^CTD^-S) by fluorescence anisotropy. Binding of RIF1^CTD^-L and RIF1^CTD^-S to an antiparallel G4 substrate was indistinguishable (Sup. Fig. 6G). Hence, we conclude that these canonical measurements of RIF1 function are not significantly impacted by the S/K cassette.

### Isoform proteomics suggests RIF1 functional differences in genome protection and DSBR

We carried out quantitative proteomic analysis of GFP-RIF1-L and GFP-RIF1-S stably expressed in *RIF1^-/-^* U-2 OS cells. Because relevant RIF1 interactions are likely to occur in the context of chromatin, we adapted the crosslinking-based RIME (rapid immunoprecipitation mass spectrometry of endogenous proteins) in conjunction with label-free quantitative LC-MS/MS (41) (Fig. 5A). We combined data from two technical replicates of three independent RIME crosslinking experiments to identify proteins significantly enriched in α-GFP-RIF1-L and/or α-GFP-RIF1-S immunoprecipitates (IPs) versus α-GFP IPs (Sup. Table 2). Using an FDR of <.05, 451 proteins were significantly enriched in GFP-RIF1-L. A total of 293 proteins were identified in GFP-RIF1-S IPs, of which, 248 were also identified for GFP-RIF1-L (Fig. 5B). The reduced number of interactants for GFP-RIF1-S may reflect reduced chromatin association (see below) or its slightly lower expression in *RIF1^-/-^*U-2 OS cells (Sup. Fig. 6A).

**Figure 5.**
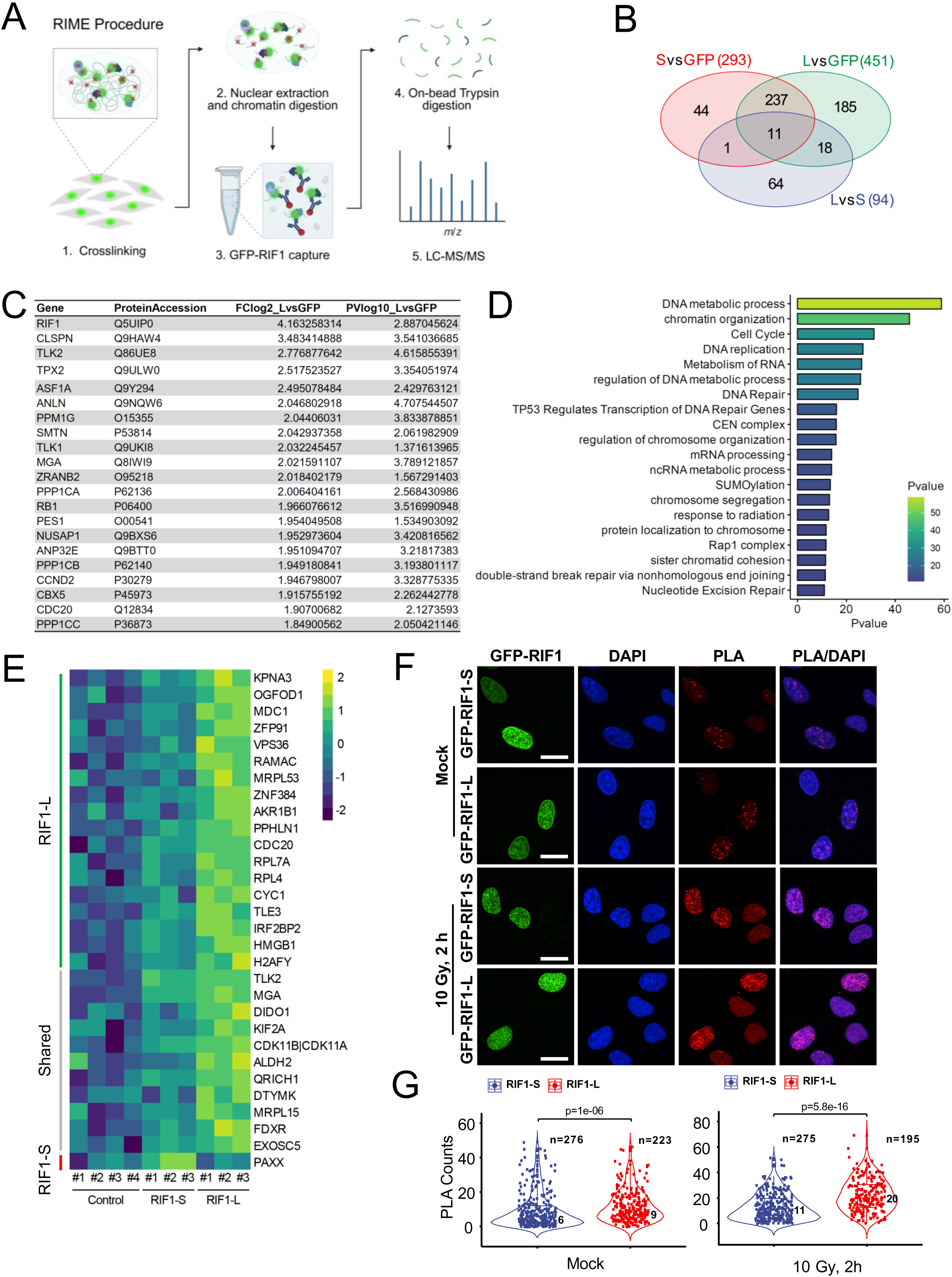
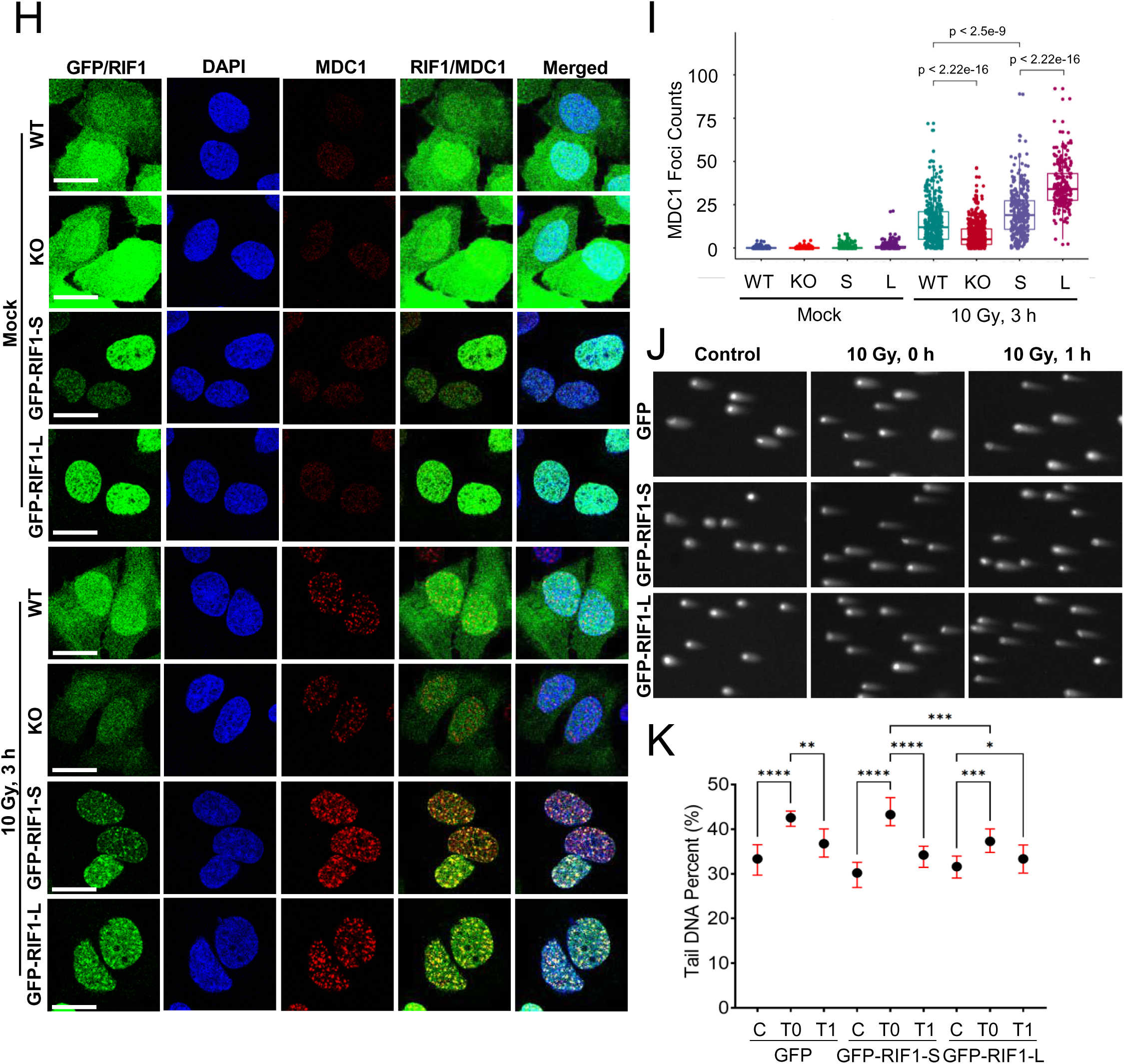
Chromatin proteomic analysis of RIF1 isoforms showed isoform-specific interactome. (A) Flow chart of the RIF1 RIME procedure. (B) Venn diagram showing proteins significantly enriched in GFP-RIF1-S (red) and GFP-RIF1-L (green) IPs (relative to GFP controls) and proteins differentially enriched between GFP-RIF1-S and GFP-RIF1-L IPs (blue). (C) Top 20 enriched proteins common to GFP-RIF1-S and GFP-RIF1-L datasets. (D) Metascape pathway analysis of the shared GFP-RIF1-S/L interactome. (E) Heat map representation of 30 proteins showing differential enrichment between GFP-RIF1-S and GFP-RIF1-L IPs. (F,G) The interaction between MDC1 and each RIF1 isoforms was evaluated by PLA using GFP, GFP-RIF1-S, and GFP-RIF1-L *RIF1^-/-^* U-2 OS cells. Cells were processed for PLA using GFP and MDC1 antibodies 2 h after exposure to 10 Gy IR or mock irradiation. The total number of cells analyzed (n), the median number of PLA foci per condition, and the p-value from Wilcox test were shown inside each violin plot. Each dot represents the PLA count number from an individual cell. Scale bar = 20 µm. (H, I) Representative images showed MDC1 foci formation enhanced by RIF1-L. *RIF1^+/+^*(WT), *RIF1^-/-^* (KO), *RIF1^-/-^*:GFP-RIF1-S, and *RIF1^-/-^*:GFP-RIF1-L U-2 OS cells were mock irradiated or exposed to 10 Gy IR followed by 3 h recovery and then fixed and stained with MDC1 antibodies. WT and KO cells were additionally stained with anti-RIF1 antibodies (Santa Cruz sc515573, 1:100). Foci analysis was performed on a minimum of 50 cells per genotype on CellProfiler. The p-values from Wilcox test were shown in the plot. Each dot represents the number of MDC1 foci from an evaluated cell. (J) Representative images from neutral comet assays of GFP, GFP-RIF1-S and GFP-RIF1-L *RIF1^-/-^*U-2 OS cells. Nucleoids from lysed cells which were exposed to 10 Gy irradiation were collected at time = 0 h or 1 h post-irradiation and electrophoresed in neutral TBE buffer. (K) Quantification of the mean comets’ tail DNA percent ± standard error pooled from three biological replicates, each with at least 50 comets scored. C = Control/mock irradiated cells; T0 = 10 Gy irradiated cells collected at 0 h post-irradiation; T1 = 10 Gy irradiated cells collected at 1 h post-irradiation. Total comet number ranging from 343 to 439 for each group (*. p ≤ 0.05; **. p ≤ 0.01; ***. p ≤ 0.001; ****. p ≤ 0.0001 from one-way Brown-Forsythe and Welch ANOVA with Games-Howell’s multiple comparisons test).

The dataset of shared RIF1-interacting proteins contained known RIF1 interactors, including PP1, TLK2, ASF1, and 53BP1 (42–45) (Fig. 5C) as well as factors not previously reported to associate with RIF1. Outside of RIF1 itself, the most highly enriched protein in RIF1-L and RIF1-S IPs was Claspin, an adaptor protein that facilitates ATR-dependent activation of the effector kinase CHK1 in response to DNA replication inhibition (46, 47). Other novel RIF1-associated proteins identified in both RIF1-S and RIF1-L IPs include TLK2, a protein kinase implicated in nucleosome assembly, DNA replication and DNA repair; TOP2A, and several proteins involved in mitosis, including TPX2, KIF4A, KIF23, and CENPF (48–52); the histone chaperone ANP32E (53); the telomerase-associated pescadillo ribosomal biogenesis factor 1 (PES1); (54) and the mitotic checkpoint regulator CDC20 (55, 56) (Fig. 5C). Metascape analysis identified DNA metabolic process, chromatin organization, cell cycle, DNA replication, and DNA repair as overrepresented gene functional groups in RIF1-L/S chromatin proteomes (Fig. 5D).

A comparison of the GFP-RIF1-L and GFP-RIF1-S RIME datasets yielded a total of 94 differentially enriched proteins (Fig. 5B); however, 64 of these were excluded from further analysis because they were not significantly enriched in either GFP-RIF1-L or GFP-RIF1-S IPs relative to GFP controls. Of the remaining 30 proteins, 11 were detected in both GFP-RIF1-L and GFP-RIF1- S, but were more enriched in GFP-RIF1-L; 18 proteins were selectively enriched in GFP-RIF1-L IPs; and 1 protein, the NHEJ regulator PAXX (57), was selectively enriched in GFP-RIF1-S IPs (Fig. 5E).

The two proteins showing the greatest fold-change difference between GFP-RIF1-L and GFP- RIF1-S IPs were the nuclear import receptor karyopherin A 3 (KPNA3) and mediator of DNA damage checkpoint 1 (MDC1) (Fig. 5E, Sup. Table 2). The apparent ∼4-fold enrichment of MDC1 in RIF1-L IPs is consistent with a study by Gupta *et al.* that identified RIF1 peptides in MDC1 proximity labeling studies (9), and was particularly interesting given that MDC1 recruits RNF8 and consequently 53BP1 and RIF1 to the sites of DNA damage (5, 49–51, 58–60). In support of the RIME-MS analysis, a proximity ligation assay (PLA) revealed that endogenous MDC1 was associated with both RIF1-L and RIF1-S (Fig. 5F) and that the number of PLA foci was significantly greater in GFP-RIF1-L versus GFP-RIF1-S U-2 OS cells, and this interaction was further strengthened upon irradiation (Fig. 5F,G). Given these findings, we evaluated IR-induced MDC1 focus formation in *RIF1^+/+^*, *RIF1^-/^*^-^, GFP-RIF1-L, and GFP-RIF1-S U-2 OS cells. The number of MDC1 foci was significantly reduced *in RIF1^-/-^* cells relative to *RIF1^+/+^*cells 3 h after exposure to IR, suggesting RIF1 enhances stable MDC1 recruitment (Fig. 5H,I). Furthermore, MDC1 foci were more abundant in GFP-RIF1-L U-2 OS cells versus GFP-RIF1-S U-2 OS cells, suggesting that RIF1-L amplifies MDC1 accumulation at DSBs.

The selective enrichment of PAXX in GFP-RIF1-S RIME IPs suggests a direct link between RIF1- S and NHEJ. Because PAXX participates in NHEJ through interaction with Ku70 and promoting DSB end synapsis (57, 61–66), we tested RIF1-L and RIF1-S cells for global DSBR activity using a neutral comet assay, which primarily measures the liberation of double-stranded DNA fragments from permeabilized nuclei. Initial levels of DSBs were significantly lower in GFP-RIF1-L cells versus GFP-RIF1-S and GFP U-2 OS cells immediately following exposure to 10 Gy IR at T0, suggesting a potential role in genome protection for RIF1-L (Fig. 5J,K). The levels of residual DSBs an hour post-IR (T1) were comparable between GFP-RIF1-L and GFP-RIF1-S cells but slightly elevated in GFP controls (Fig. 5J,K). Thus, while GFP-RIF1-S cells were more prone to IR-induced DSB, the steeper slope between T0 and T1 suggests an enhanced rate of DSBR in GFP-RIF1-S versus GFP-RIF1-L cells.

### CTD phosphorylation diminishes RIF1 chromatin and MDC1 association

Orbitrap MS identified several phosphorylation sites in GFP-RIF1^CTD^-L, including S2205, which is located in the PP1 binding site; S2260 and S2265, which are located in Ex32-encoded S/K cassette (Fig. 6A); and S2348, which lies within CR2. We generated phospho-specific antibodies against a peptide dually phosphorylated on S2260 and S2265 (α-RIF1-pS2260/65) and validated the site-specificity of the antibody in Western blotting experiments using GFP-RIF1-L^1A^ and GFP- RIF1-L^2A^ mutants with Ala mutation at site(s) S2260 and S2260/65. GFP-RIF1-L^1A^ reduced α- RIF1-pS2260/65 recognition dramatically while GFP-RIF1-L^2A^ completely abolished recognition by this antibody (Fig. 6B). Using Nocodazole-synchronized U-2 OS cells, we found that RIF1- pS2260/65 phosphorylation was maximal in mitosis and rapidly extinguished following mitotic exit (Fig. 6C).

**Figure 6.**
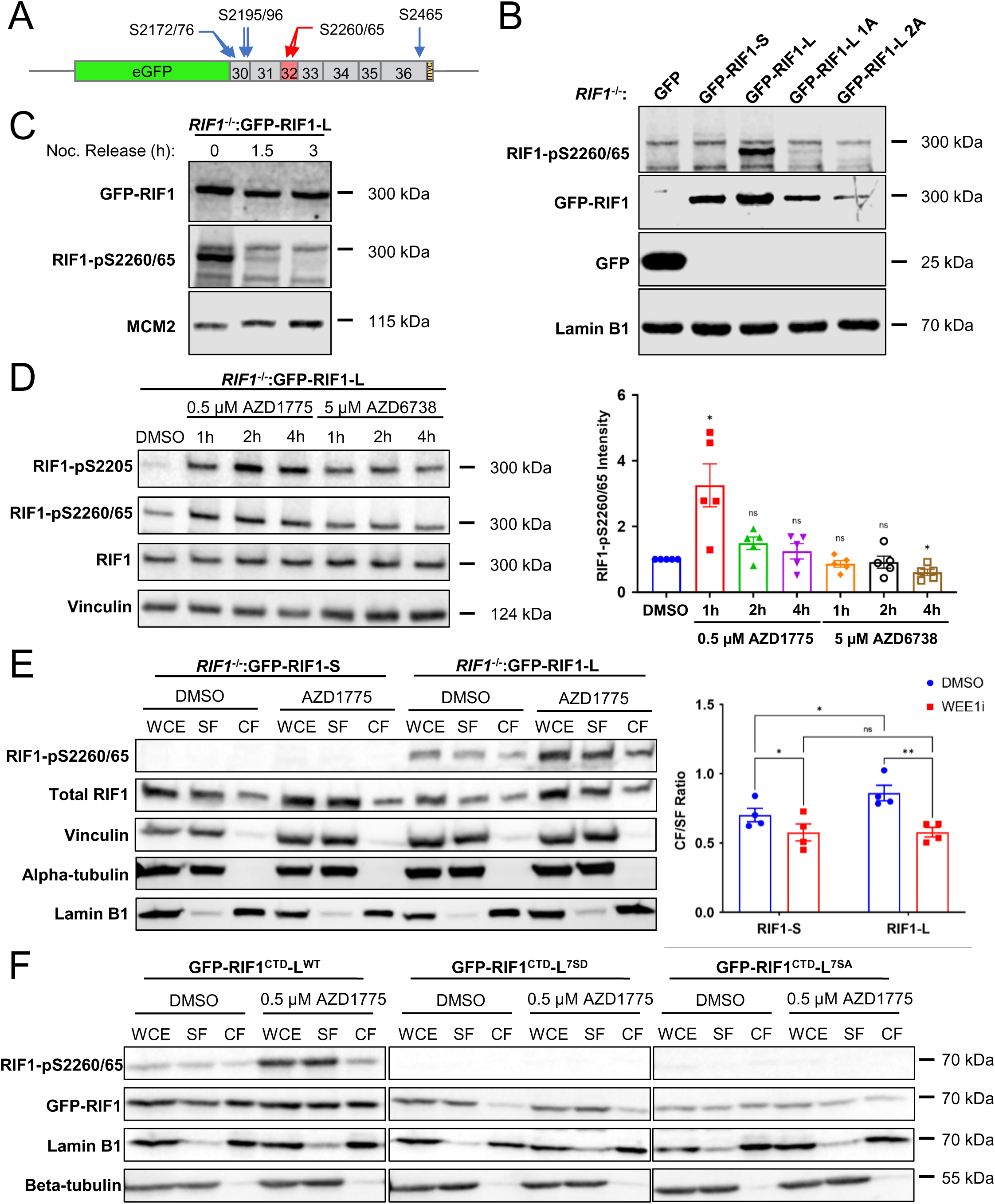
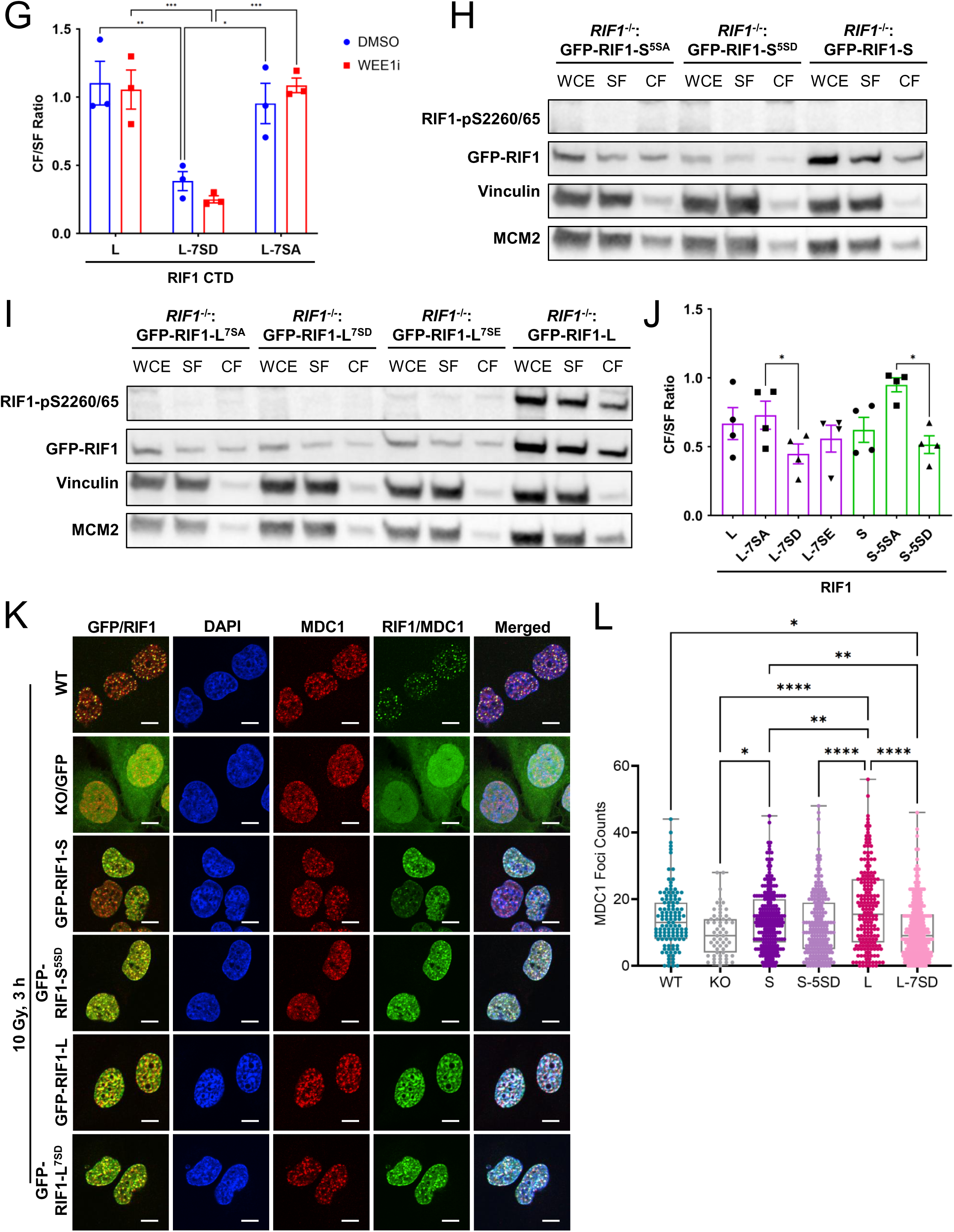
RIF1 phosphorylation on S2260 and S2265 during mitosis and in response to WEE1 inhibitors decreases its chromatin association. (A) Schematic of GFP-RIF1^CTD^-L showing five UniProt-annotated phospho-Ser residues (blue arrows) and S2260/65 (red arrows). These selected Ser residues were subsequently mutated to Asp or Ala. (B) Specificity of α-RIF1- pS2260/65 antisera was tested with lysates from *RIF1^-/-^* U-2 OS cells expressing GFP or the indicated RIF1 alleles. RIF1-L^1A^ and RIF1-L^2A^ alleles harbor Ala mutations at S2260 and S2260/S2265, respectively. (C) *RIF1^-/-^*:GFP-RIF1-L U-2 OS cells were synchronized in prometaphase with nocodazole (Noc) and released into Noc-free media for the indicated lengths of time. Note rapid reduction in RIF1-pS2260/65 levels following Noc release. (D, left panel) *RIF1^-/-^*:GFP-RIF1-L U-2 OS cells were treated with WEE1 inhibitor (AZD1775) or ATR inhibitor (AZD6738) for the indicated timepoint prior to immunoblotting analysis with the indicated antibodies. (D, right panel) Quantification of left panel based on densitometry. Bar height represents mean intensity of RIF1-pS2260/65 of each treatment normalized to the baseline phosphorylation level in DMSO control ± standard error. Each dot represents an individual biological replicate, N = 5 (ns. not significant; *. p ≤ 0.05 from two-tailed one sample t-test). (E, left panel) Representative Western blot showing the chromatin fractionation patterns for GFP- RIF1-L and GFP-RIF1-S cells treated with DMSO or 0.5 µM AZD1775 for 1 h. Whole-cell extract (WCE), chromatin fraction (CF), and soluble fraction (SF) were resolved by SDS-PAGE and blotted with the indicated antibodies. (E, right panel) Quantification of the mean chromatin/soluble fraction (CF/SF) ratio of total RIF1-L/RIF1-S ± standard error based on densitometry. Each dot represents an individual biological replicate, N = 4 (ns. not significant; *. p ≤ 0.05; **. p ≤ 0.01 from two-way ANOVA with uncorrected Fisher’s LSD test). (F, G) Chromatin fractionation patterns of GFP-RIF1^CTD^-L and its serine mutants. The indicated GFP-RIF1^CTD^-L constructs were transiently expressed in U-2 OS cells treated with DMSO or 0.5 AZD1775 for an hour. Note that phospho- GFP-RIF1^CTD^-L^WT^ and phosphomimetic GFP-RIF1^CTD^-L^7SD^ were highly enriched in the SF, while GFP-RIF1^CTD^-L^7SA^ with abolished phosphorylation sites was not. (G) Quantification of the mean CF/SF ratio of GFP-RIF1 in (F) ± standard error based on densitometry. Each dot represents an individual biological replicate, N = 3 (*. p ≤ 0.05; **. p ≤ 0.01; ***. p ≤ 0.001 from two-way ANOVA with Tukey’s multiple comparisons test). (H, I) Chromatin fractionation patterns of full-length GFP- RIF1 isoforms and its serine mutants. *RIF1^-/-^* U-2 OS cells expressing the indicated GFP-RIF1-S (H) and GFP-RIF1-L (I) constructs were subjected to chromatin fractionation. (J) Quantification of the mean CF/SF ratio of GFP-RIF1 in (H) and (I) ± standard error based on densitometry. Each dot represents an individual biological replicate, N = 4 (*. p ≤ 0.05; from repeated measures one- way ANOVA with Geisser-Greenhouse correction and Tukey’s multiple comparisons test). (K) Representative MDC1 foci images of *RIF1^+/+^* (WT), *RIF1^-/-^*:GFP (KO/GFP), *RIF1^-/-^*:GFP-RIF1-S, *RIF1^-/-^*:GFP-RIF1-S^5SD^, *RIF1^-/-^*:GFP-RIF1-L, and *RIF1^-/-^*:GFP-RIF1-L^7SD^ U-2 OS cells after 10 Gy irradiation followed by 3 h recovery before being stained with MDC1 antibodies (Sigma HPA006915, 1:500). WT cells were additionally stained with anti-RIF1 antibodies (Santa Cruz sc515573, 1:100). Scale bar = 10 µm. (L) Box plots showing the quantification of MDC1 foci counts per genotype by ImageJ macro Foci_Analyzer_1_5. Each dot represents the number of MDC1 foci from an evaluated cell, total cells scored ranging from 59 to 277 for each genotype (*. p ≤ 0.05; **. p ≤ 0.01; ****. p ≤ 0.0001 from one-way ANOVA with Tukey’s multiple comparisons test).

S2260 and S2265 residues occur in a Ser-Pro dipeptide motif that is a consensus for the mitotic cyclin-dependent kinase 1 (CDK1) and related kinases. A recent study suggested that, under conditions of ATR inhibition, CDK1-dependent phosphorylation of RIF1 on S2205 diminished PP1 binding, leading to increased phosphorylation stoichiometry of CDK2 and CDC7 substrates and elevated rates of origin firing (67, 68). To determine whether S2260 and S2265 phosphorylation exhibit a similar phosphorylation profile, we measured RIF1-pS2260/65 levels in GFP-RIF1-L U- 2 OS cells cultured in the presence of a WEE1 inhibitor (AZD1775) or an ATR inhibitor (AZD6738). AZD1775 treatment significantly increased RIF1-pS2260/65 level an hour after treatment, while ATR inhibitors had little effect (Fig. 6D). Thus, RIF1-S2205 and RIF1-S2260/65 phosphorylation sites are co-phosphorylated under conditions of aberrant CDK1 activation.

The basic nature of the S/K cassette suggested it may play a role in nuclear localization, DNA binding, and/or chromatin association. While RIF1-L and RIF1-S exhibited comparable nuclear localization and binding to G-quadraplex DNA substrates (Sup. Fig. 6B,G), chromatin fractionation of full-length GFP-RIF1-L and GFP-RIF1-S suggested that RIF1-L has significantly higher chromatin binding affinity relative to RIF1-S (Fig. 6E, blue bars). WEE1 inhibition using AZD1775 increased the proportion of S2260/65-phosphorylated RIF1-L (RIF1-pS2260/65) and decreased the chromatin association of both RIF1-L and RIF1-S (Fig. 6E, red bars).

To further explore the relationship between RIF1 phosphorylation and its chromatin association, we compared the chromatin association profiles of GFP-RIF1^CTD^-L^WT^ to that of GFP-RIF1^CTD^-L^7SA^, and GFP-RIF1^CTD^-L^7SD^ mutants harboring seven Ser-Ala (SA) or Ser-Asp (SD) mutations at UniProt-annotated CDK1 phosphorylation sites, including S2260 and S2265, within the S/K cassette (Fig. 6A). Similar to what was observed using full-length RIF1-L, the phosphorylation of RIF1^CTD^-L^WT^ on S2260/65 was significantly increased in response to WEE1 inhibition with AZD1775 (Fig. 6F, left panel). In addition, the RIF1-pS2260/65 signal was strongly enriched in the soluble fraction (SF) relative to the chromatin fraction (CF), suggesting that phosphorylation reduces RIF1^CTD^ chromatin-binding affinity. Consistent with this, the phosphomimetic RIF1^CTD^- L^7SD^ mutant exhibited a significant reduction in chromatin association even without AZD1775 treatment (Fig. 6F, center panel and Fig. 6G). While the RIF1^CTD^-L^7SA^ mutant exhibited lower expression, its chromatin binding profile was similar to RIF1^CTD^-L^WT^ (Fig. 6F, right panel and Fig. 6G). As expected, Western blotting with RIF1-pS2260/65 antibodies did not yield a signal in U-2 OS cells expressing either RIF1^CTD^-L^7SD^ or RIF1^CTD^-L^7SA^. Altogether, findings with full-length RIF1 and RIF1^CTD^ fragments suggest that multiple phosphorylation sites within the RIF1 CTD, including S2260/2265 in the S/K cassette, diminish RIF1 chromatin binding in response to WEE1 inhibition.

Based on these findings, we reconstituted *RIF1^-/-^* U-2 OS cells with full-length GFP-RIF1-S^5SA^, GFP-RIF1-L^5SD^, GFP-RIF1-L^7SA^, GFP-RIF1-L^7SD^, and GFP-RIF1-L^7SE^ constructs. Consistent with what was observed with GFP-RIF1^CTD^ fragments, abolishing phosphorylation sites by SA mutations increased chromatin retention, while phosphomimetic SD and Ser-Glu (SE) mutants showed lower chromatin association (Fig. 6H-J). Since the RIF1-L-MDC1 association occurred in the context of chromatin (Fig. 5), we asked whether RIF1 CTD phosphosite mutants showed any defect in MDC1 focus formation. Consistent with findings in Fig. 5I, GFP-RIF1-L rescued the MDC1 focus formation defect of *RIF1^-/-^* cells to a greater extent than GFP-RIF1-S U-2 OS cells (Fig. 6K,L). The GFP-RIF1-L^7SD^ mutant showed significantly fewer MDC1 foci versus GFP- RIF1^WT^-L. From this we surmise that phosphorylation of the S/K cassette and flanking CTD diminishes RIF1 association with MDC1 on chromatin.

### The S/K cassette regulates RIF1 phase separation

Using the DISOPRED 3.1 disorder prediction tool (69), we found that the presence of the S/K cassette reduced disorder of the region roughly spanning amino acids ∼2240 to 2280 in the RIF1 CTD (Fig. 7A). In transient transfection assays, GFP-RIF1^CTD^-L and GFP-RIF1^CTD^-S formed spherical shells in the nuclei of U-2 OS cells that ranged from single shells to complex arrangements containing multiple chambers (Fig. 7B,C). In contrast, GFP-RIF1^CTD^-L^ΔNLS^ with a deletion of the nuclear localization signal (NLS) showed diffused GFP signal in the cytoplasm (Fig. 7B). Three-dimensional reconstruction revealed RIF1 nuclear shells to be oblong spheroids with a hollow central core (Sup. Video 1). Because they closely resembled the birefringent “anisosomes” formed by the nuclear RNA-binding protein TDP-43 (70), we have adopted the anisosome nomenclature to describe RIF1^CTD^ nuclear assemblies.

**Figure 7.**
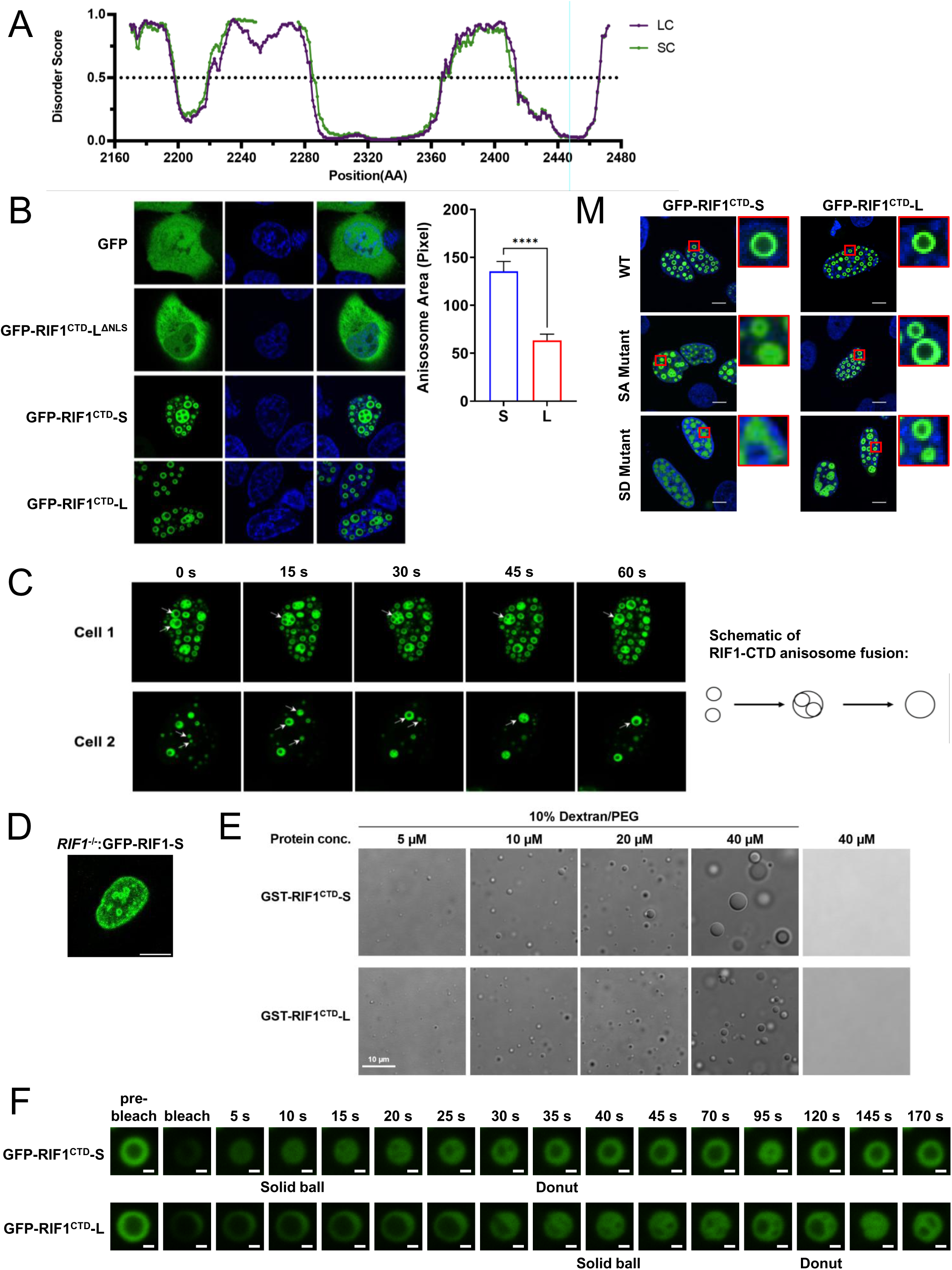
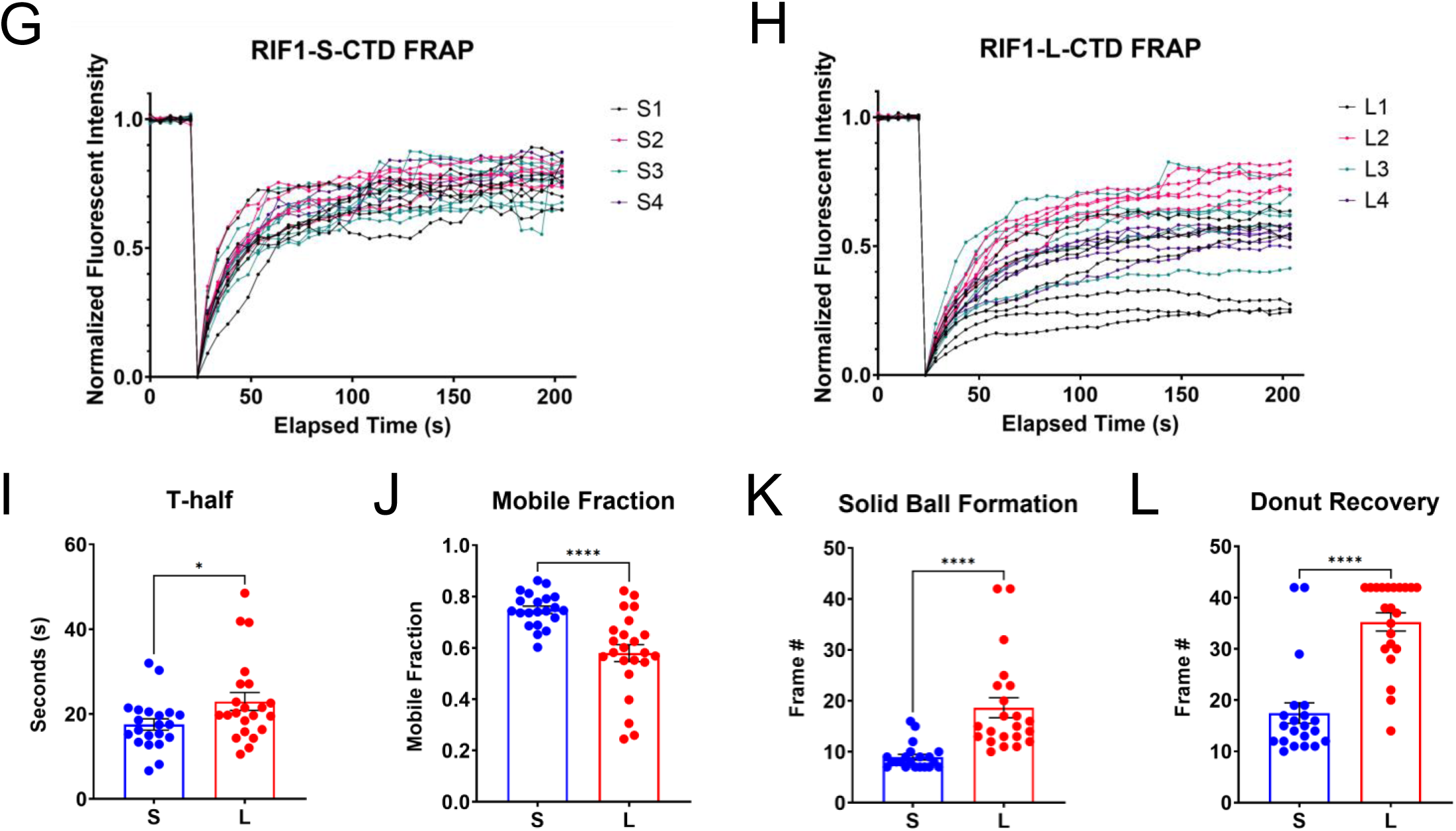
The S/K cassette stabilizes phase separation of the RIF1 CTD. (A) Predictive disorder score of GFP-RIF1^CTD^-S and GFP-RIF1^CTD^-L showed that the presence of S/K cassette (2250 – 2275 aa) decreases the disorder of RIF1 CTD. (B, left panel) Representative examples of RIF1 CTD anisosomes. U-2 OS cells were transiently transfected with GFP, GFP-RIF1^CTD^-L^ΔNLS^, GFP-RIF1^CTD^-S or GFP-RIF1^CTD^-L and subjected to live cell imaging after Hoechst 33342 staining (blue). (B, right panel) Quantification of mean anisosome area ± standard error by customized ImageJ script. N = 586 for GFP-RIF1^CTD^-S; N = 396 for GFP-RIF1^CTD^-L (***. p ≤ 0.0001 from unpaired two-tailed t-test assuming equal standard deviation). (C) Timelapse images and the schematic showing the fusion of GFP-RIF1^CTD^ anisosomes in 60 seconds. Single anisosomes and the subsequently fused multi-chambers anisosomes were marked with white arrows. (D) Full-length GFP-RIF1-S occasionally forms anisosome-like structure in Dox-inducible *RIF1^-/-^*U-2 OS cells. Scale bar = 10 µm. (E) *In vitro* phase separation assays showed concentration (10-40 µM) dependent LLPS droplet formation of purified GST-RIF1^CTD^-S and GST-RIF1^CTD^-L proteins. Scale bar = 10 µm. (F) Fluorescence recovery after photobleaching (FRAP) montage for representative GFP-RIF1^CTD^-S and GFP-RIF1^CTD^-L anisosomes in U-2 OS cells. Scale bar = 1 µm. (G, H) FRAP recovery curves of the bleached anisosomes for GFP-RIF1^CTD^-S and GFP-RIF1^CTD^-L over a time course of three minutes. Four biological replicates were carried out, each with at least three technical replicates. Each recovery curve was color-coded according to the biological replicate number (S1-4 or L1-4) in each plot. (I, J) Each recovery curve from (G) and (H) was fitted by single exponential equation to estimate the t-half value of recovery and mobile fraction of the anisosome. Bar height represents mean ± standard error of the pooled data. Each dot represents score from an anisosome, N = 21 for GFP-RIF1^CTD^-S and N = 22 for GFP-RIF1^CTD^-L (*. p ≤ 0.05; ****. p ≤ 0.0001 from unpaired two-tailed t-tests with Welch’s correction which do not assume equal standard deviation in populations). (K, L) Manual tabulation of the recovery frame where the two distinct stages – “solid ball” and “donut”, as indicated in (F) – reappeared after photobleaching was done. Bar height represents mean ± standard error of the pooled data. Each dot represents score from an anisosome, N = 21 for GFP-RIF1^CTD^-S and N = 22 for GFP-RIF1^CTD^- L (*. p ≤ 0.05; ****. p ≤ 0.0001 from unpaired two-tailed t-tests with Welch’s correction). (M) Representative images showed that serine to alanine (SA) and serine to aspartic acid (SD) mutations impede anisosome formation to a greater extent in GFP-RIF1^CTD^-S expressing cells. Red boxes showed enlargement of the anisosomes of interest. Scale bar = 10 µm.

Time lapse imaging revealed RIF1^CTD^ nuclear anisosomes to be dynamic structures that frequently fused to form larger single- or multi-chamber structures (Fig. 7C, Sup. Video 2). In addition, GFP-RIF1^CTD^ anisosomes underwent spontaneous cycles of closure and reopening (Sup. Video 2). While both GFP-RIF1^CTD^-S and GFP-RIF1^CTD^-L formed anisosomes, they exhibited different properties. GFP-RIF1^CTD^-S anisosomes exhibited an increased rate of fusion events versus GFP-RIF1^CTD^-L anisosomes to form larger structures (Fig. 7B), while GFP-RIF1^CTD^-L anisosomes occasionally formed nested structures—rarely observed for GFP-RIF1^CTD^-S—in which smaller anisosomes were enclosed within a larger assembly (Sup. Video 3). These nested anisosomes may be fusion intermediates and were therefore observed more frequently in the less dynamic GFP-RIF1^CTD^-L which fused slower. RIF1^CTD^ nuclear anisosomes were also observed in transiently transfected HeLa and HEK293T cells as well as U-2 OS cells expressing Dox-inducible GFP-RIF1^CTD^-S and GFP-RIF1^CTD^-L (not shown). Full-length GFP-tagged RIF1 protein, on rare occasion, can also form anisosome-like droplets as shown in Fig. 7D.

The assembly of GFP-RIF1^CTD^ into anisosomes suggested the CTD undergoes liquid-liquid phase separation (LLPS). To test this, we incubated purified, GST-tagged RIF1^CTD^-S and RIF1^CTD^-L with 10% Dextran/PEG. Both GST-RIF1^CTD^-S and GST-RIF1^CTD^-L underwent concentration-dependent LLPS, however, RIF1^CTD^-S formed significantly larger droplets compared to RIF1^CTD^- L with increasing RIF1^CTD^ concentration (Fig. 7E), possibly reflecting increased rates of droplet fusion seen in transient transfection assays (Fig. 7C).

To better evaluate the dynamics of RIF1^CTD^ anisosomes, we performed fluorescence recovery after photobleaching (FRAP) experiments in which GFP-RIF1^CTD^-S and GFP-RIF1^CTD^-L anisosomes were allowed to recover after photobleaching. Both GFP-RIF1^CTD^-S and GFP- RIF1^CTD^-L anisosomes rapidly disintegrated following photobleaching, losing their donut-like character with laser exposure before reassembling over the course of 3 min (Fig. 7F, time course/montage, Sup. Video 4-5). The recovery curve of each bleached anisosome was then plotted from the average fluorescent intensity of the bleached region over time (Fig. 7G,H). GFP- RIF1^CTD^-S anisosomes showed a small but significantly lower t-half value and a larger mobile fraction estimated from the recovery curves of RIF1^CTD^-S and RIF1^CTD^-L anisosomes (Fig. 7I,J). Due to the larger heterogeneity within RIF1^CTD^-L anisosome population (Fig. 7H) and the difficulty in measuring anisosome morphological recovery based on average fluorescent intensity, we also tabulated the frame numbers in which a bleached anisosome formed either a solid ball intermediate or a fully recovered “donut” as indicated in Fig. 7F. The formation of both structures after photobleaching was significantly delayed for GFP-RIF1^CTD^-L versus GFP-RIF1^CTD^-S, suggesting that GFP-RIF1^CTD^-L anisosomes are intrinsically more stable than GFP-RIF1^CTD^-S anisosomes (Fig. 7K,L). Although the reason for GFP-RIF1^CTD^-L anisosome recovery heterogeneity is unclear, it is conceivable that the expanded interactome for RIF1^CTD^-L or posttranslational modification of the S/K cassette influences its phase separation dynamics.

To investigate the potential role of phosphorylation in RIF1 CTD phase separation, we transfected U-2 OS cells with GFP-RIF1^CTD^-L and GFP-RIF1^CTD^-S CTDs harboring the corresponding 7SA/SD and 5SA/SD mutations (Fig. 6A). In contrast to the well-demarcated anisosomes formed by wild-type GFP-RIF1^CTD^-S, GFP-RIF1^CTD^-S^5SA^ formed irregularly shaped anisosomes that typically featured a narrow central cavity and thick outer shell. Loss of anisosomal character was even more pronounced for RIF1^CTD^-S^5SD^, which exclusively formed irregular nuclear aggregates (Fig. 7M). While the corresponding GFP-RIF1^CTD^-L^7SA^ and GFP- RIF1^CTD^-L^7SD^ mutants also exhibited anisosome morphology defects; the changes were less dramatic than those seen in GFP- RIF1^CTD^-S^5SA^ and GFP-RIF1^CTD^-S^5SD^. This was most pronounced for the SD mutations that completely disrupted GFP-RIF1^CTD^-S anisosomes but only partially inhibited GFP-RIF1^CTD^-L anisosome formation. The fact that the presence of S/K cassette reduced the impact of disruptive serine mutations on anisosome formation supports the conclusion that this motif stabilized phase separation of the RIF1 CTD.

## DISCUSSION

RIF1 fulfills evolutionarily conserved and remarkably diverse roles in maintaining genome stability. Here we show that DNA damage- and cell cycle-regulated alternative splicing contributes to RIF1 functional diversity by altering the biochemical properties of the RIF1 CTD. Consequently, RIF1- L and RIF1-S exhibit measurable differences in chromatin association, phase separation, and association with DNA repair factors. We propose that these functional differences allow for optimized RIF1 function in response to genotoxic stress and in different cellular contexts.

The AS of RIF1-Ex32 in mammalian cells was regulated by no less than six RBPs, including two positive regulators (SRSF1 and snRNP70) and four negative regulators (PTBP1, SRSF3, SRSF7, and RBM28) which functionally interact with one another and may act cumulatively or synergistically to promote Ex32 skipping. The central role for SRSF1 as a positive regulator of Ex32 inclusion was supported by its association with RIF1-L mRNA, which was reduced in *RIF1- L^-/-^* cells that harbor a point insertion (denoted by ^) in a CCCAGGA^T SRSF1 recognition motif (Fig. 3A,4F). RIF1-Ex32 was also identified as an SRSF1-regulated cassette exon in developing mouse epithelia by Yu *et al.* (34). Although snRNP70 binding was not detected (possibly due to poor antibody performance in RNA-IP), it is often found within the same alternative splicing complex as SRSF1 where it enhances recognition of the 5’ splice site (34, 71). Cooperative activities of SRSF1 and snRNP70 may promote RIF1-Ex32 inclusion in actively dividing progenitors and S/G_2_-phase cells where RIF1-L levels are high (Fig. 1I,J).

DNA damage promoted Ex32 skipping, leading to increased production of RIF1-S mRNA and protein (Fig. 1). Inhibition of Ex32 splice-in could be caused by inhibition of positive splicing regulators (e.g. SRSF1), activation of negative splicing regulators (e.g. SRSF3, SRSF7, and PTBP1), or a combination of mechanisms. Interestingly, while SRSF1, SRSF3, and SRSF7 were all induced by DNA damage (Fig. 4L,M), only SRSF3 and SRSF7 showed increased occupancy over RIF1 transcripts in CLM-treated cells (Fig. 4I-K). This suggests that the inhibitory effects of SRSF3 and SRSF7 binding are dominant to the positive effects of SRSF1 binding as it pertains to Ex32 inclusion in the setting of DNA damage. Because one primer set detected a modest reduction in SRSF1-binding in CLM-treated cells (Fig. 4I), it is possible that SRSF3 and SRSF7 directly compete with SRSF1 for binding to sequences in the 3’ end of Ex32, which is consistent with the SRSF1 binding motif identified in our *RIF1^-/-^*cells (Fig. 3A). Perhaps relevant to these data, previous work has shown that SRSF3 levels are maintained within a narrow range through negative autoregulation, as well as positively regulated by SRSF1 and SRSF2 as reviewed in Ref.(72) (73–75). Hence, increased SRSF1 expression following DNA damage could activate SRSF3, thereby contributing to the RIF1-L to RIF1-S isoform switch. On the other hand, PTBP1 impairs SRSF3 auto-regulation which likely explains the overexpression of both in primary cancers which predominantly express RIF1-S (34, 71, 72, 76) (Fig. 2, Sup. Fig. 5B). It is perhaps noteworthy that the 3’ end of RIF1-In31 contains a conserved pyrimidine-rich tract (TTTTTTTCTCTCCTTTCTTCT) that may mediate PTBP1 binding. Regardless of the exact mechanism, our findings suggest that the rates of RIF1-Ex32 inclusion are determined by the competitive balance between positive (e.g. SRSF1) and negative (SRSF3, SRSF7, and PTBP1) regulators, whose abundance and activities are regulated by DNA damage and the cell cycle. In the G1 phase and in the scenario of prolonged DNA damage, the inhibitory splicing factors prevail, leading to increased levels of RIF1-S. Deciphering potential cooperative interactions between these factors and understanding how their activities are affected by DNA damage will be the subject of future studies.

RIF1 chromatin proteomics identified both known and novel RIF1 interactants and pointed toward MDC1 as a potential target of regulation by RIF1-L (Fig. 5E-I). Because MDC1 is also a chromatin-associated protein (77), its increased abundance in RIF1-L IPs could be due to the enhanced chromatin association of RIF1-L versus RIF1-S (Fig. 6E). Alternatively, the S/K cassette could mediate direct interaction with MDC1. While the functional significance of the enhanced RIF1-L- MDC1 interaction is not fully understood, our findings suggest that RIF1-L sustains MDC1 accumulation at IR-induced foci and phosphorylation of its CTD reduces this property (Fig. 5H,I, Fig. 6K-L). This finding was somewhat unexpected given that RIF1 is recruited to IR-induced foci downstream of MDC1 through direct interaction with phosphorylated 53BP1 (28). How RIF1 promotes MDC1 focus formation is unclear. One speculation is that oligomerization of chromatin-bound RIF1-L participates in the recently described clustering of topologically associating domains (TADs) that have incurred DNA damage (78), leading to local enrichment of MDC1. We also note that RIF1-L was reported to enhance 53BP1 nuclear body assembly in G_1_-phase cells (79). Although not tested here, such a function is conceptually consistent with the enhancement of DNA damage focus formation seen in our studies and might be akin to RIF1 function in the organization of TADs with shared replication timing (25). Further experiments are needed to establish mechanisms and functional consequences of the RIF1-L-MDC1 interaction.

Finally, while the functional significance is not known, the only protein that showed enhanced interaction with RIF1-S was the auxiliary NHEJ factor, PAXX. RIF1-S association with PAXX suggests the intriguing possibility that, in addition to its roles in suppressing HDR, RIF1 may also contribute an isoform-specific role in mediating NHEJ directly (Fig. 5E). Although this hypothesis remains to be tested, RIF1-L and RIF1-S U-2 OS cells exhibited distinct DSBR profiles post-IR in neutral comet assays that may be compatible with differences in DSBR rate. Specifically, even though RIF1-S expressing cells were more susceptible to IR-induced DSBs, the breaks were repaired significantly faster compared to *RIF1^-/-^* and RIF1-L expressing cells (Fig. 5J,K). Although the precise nature of the RIF1-S-PAXX interaction awaits further study, we note that RIF1-S mRNA is induced by DNA damage and most abundant in G_1_ phase when NHEJ is the predominant DSBR pathway (Fig. 1I,J).

The basic S/K cassette strengthened the chromatin binding of RIF1-L which was diminished through multisite phosphorylation of the CTD—including at least two sites within the S/K cassette—in response to WEE1i (Fig. 6D-F). Previous work showed that WEE1 inhibitors induced CDK1-dependent phosphorylation of RIF1 on S2205, leading to PP1 dissociation, MCM4 hyperphosphorylation, and activation of dormant replication origins (67). Given that phosphorylation of S2260/65 and flanking CDK1 site occurs concomitant with S2205 phosphorylation (Fig. 6D), we propose that chromatin eviction of RIF1 contributes to the loss of origin suppression seen in cells treated with WEE1 inhibitors (67).

The RIF1 CTD undergoes LLPS *in vitro* and phase-separates into anisosome-like structures in intact cells (70) (Fig. 7B). While both GFP-RIF1^CTD^-S and GFP-RIF1^CTD^-L formed anisosomes, they exhibited distinct characteristics. GFP-RIF1^CTD^-S anisosomes were larger, more dynamic, fusion prone (Fig. 7C), and exhibited faster FRAP recovery times with larger mobile fraction as a more homogenous species compared to GFP-RIF1^CTD^-L anisosomes (Fig. 7F-L). GFP-RIF-L^CTD^ was less susceptible to disruption by phosphomimetic amino acid substitutions (Fig. 7M), which are well known to disrupt phase separation of Fused in sarcoma (FUS) and other proteins that undergo LLPS (25, 80). This finding is consistent with studies showing that Ser-rich motifs can function as spacers to promote phase separation and droplet hardening (81). While full-length RIF1 also forms nuclear assemblies that may be attributed to phase-separation (Fig. 7D), it did not form discrete anisosomes, which may require supraphysiologic levels of RIF1 or may be suppressed by intramolecular folding. Nevertheless, stabilization of RIF1 LLPS by the S/K cassette may contribute to RIF1-dependent replication timing regulation, potentiation of MDC1 accumulation at IR-induced foci, 53BP1 nuclear body formation (79), and other chromatin-associated roles of RIF1.

The functional differences between RIF1-L and RIF1-S described here may be relevant to RIF1 isoform usage differences between normal and cancer cells. We note that SRSF1 was upregulated in osteosarcoma cell lines, such as U-2 OS (82) that expresses high levels of RIF1- L (Fig. 1C). In addition, the RIF1-L/S isoform ratio was consistently reduced in primary cancers of diverse origin (Fig. 2). Whether or not this isoform switch drives aspects of tumorigenesis is unclear, however, reduced expression of RIF1-L may negatively impact MDC1-mediated signaling and other chromatin-associated roles for RIF1 that may suppress neoplastic growth. Non-exclusively, RIF1-S may function as a pro-growth isoform and/or enhance DSBR through interaction with PAXX or other factors. Further delineation of RIF1 isoform-specific functions *in vivo* will illuminate these possibilities in carcinogenesis.

## EXPERIMENTAL PROCEDURES

### Cell culture and treatment

U-2 OS, HeLa, H460, and HEK293T cell lines were obtained from the American Type Culture Collection (ATCC). U-2 OS and its derivative cell lines were grown in McCoy’s 5A medium (Corning, 10-050-CV). HEK293T and HeLa cells were grown in DMEM medium (Corning, 10-013- CV). All cell lines were grown in medium supplemented with 10% fetal bovine serum (GeminiBio 900-108-500) and 1% Penicillin/Streptomycin (Corning, 30-002-CI) and incubated at 37°C in 5% CO_2_. For G_1_/S synchronization experiments, cells were treated with 2 mM thymidine for 19 h, released into thymidine-free growth media for 9 h, and then returned to thymidine-containing media for an additional 16 h (83). The cells were washed three times with PBS and then released into complete media for the indicated time periods. Calicheamicin γ1 (CLM) was prepared at a concentration of 4 µM stock solution in DMSO and used at a concentration of 2-10 ng/ml for 4-6 h. For checkpoint inhibitor treatments, U-2 OS cells were treated with 5 µM AZD6738 (ATRi) or 0.5 µM AZD1775 (WEE1i) diluted in DMSO for 1, 2 or 4 hours prior to harvesting.

### RNA extraction, RIF1 splicing assay, and quantitative PCR (qPCR)

Total RNA was extracted in TRIzol reagent (Invitrogen, 15596018) followed by cDNA synthesis by iScript™ cDNA Synthesis Kit (Bio-Rad, 1708891) or iScript™ gDNA Clear cDNA Synthesis Kit (Bio-Rad, 1725035) according to manufacturer’s protocols. Human *RIF1*-Ex32 splicing was evaluated by RT-PCR using primers located on Ex31 and Ex33 (RIF1-Ex31-F: 5’- AAGCAGGATTGGCAGATGAC-3’ and RIF1-Ex33-R: 5’-GATGTCAACTGGTGCCACAC-3’).

Positionally analogous primers located within mouse *Rif1*-Ex31 (5’- AAGCAGGATTGGCAGATGAC-3’) and Ex33 (5’-GATGTCAACTGATGCTGCAC-3) were used to analyze mouse *Rif1*-Ex32 splicing. Primers flanking Ex1a (RIF1-Ex1a-F: 5’- CGCCATCTTGGTCTAGGAGG-3’ and RIF1-Ex1a-R: 5’-ACGACTGGTCAGAGTCAGGT-3’) were used as a negative control. Beta-actin (Actin-F: 5’-TCCCTGGAGAAGAGCTACG-3’ and Actin-R: 5’-GTAGTTTCGTGGATGCCACA-3’) or GADPH (GAPDH-F: 5’- AATCCCATCACCATCTTCCA-3’ and GAPDH-R: 5’-TGGACTCCACGACGTACTCA-3’) was used as an internal control. RIF1 AS leads to the formation of RIF1-L and RIF1-S transcripts with a length difference of 78 nt which can be resolved subsequently by 2% w/v agarose gel electrophoresis (See Fig. 1B).

RIF1 isoform specific qPCR in human cells was performed using the following primers: RIF1-L (5’-GGATTGGCAGATGACATTGATAGA-3’; 5-TCCTTTGGCTGAAGTGGTATTATG-3’); RIF1-S (5’-CCTACTACACAATCTAAGATTTCA-3’; 5’-GCTCTAATGAGTTGTCCCA-3’); and total RIF1 (5’-CGCTGTGTCTGGTCTCCTT-3’; 5’GCACCGTCTATCAATGTCATCTG-3’) with iTaq Universal SYBR® Green Supermix (Bio-Rad, 1725124) according to manufacturer’s protocols.

### Gene editing and cloning

*RIF1^-/-^* and *RIF1^ΔEx32^* cells were generated by transient transfection of U-2 OS cells with pX459 vectors (v2, Addgene plasmid #62988) (84, 85) harboring two sgRNA sequences targeting RIF1-Ex2 (5’-CACCgAGTCTCCAACAGCGGCGCGA-3’ and 5’- AAACTCGCGCCGCTGTTGGAGACTc-3’) or RIF1-Ex32 (5’- CACCgATTTAGGGCTACGTGATCCT-3’ and 5’-AAACAGGATCACGTAGCCCTAAATc-3’) using jetPRIME^®^ (Sartorius, 101000046). Twenty-four hours after transfection, cells were selected for 72 h with 1 µg/ml puromycin and then diluted into 96-well-plates at an average density of 1 cell per well. Each single clone was isolated and screened for RIF1 knockout phenotype through immunostaining of ionizing radiation-induced foci (IRIF) and Western blotting with α-RIF1 and α- RIF1-L antibodies. All clones were sequenced around the sgRNA targeted sequence, and five clones (*RIF1^-/-^*: H1 and 2C5; *RIF1-L^-/-^*: A6, 2A2, and H11) were selected for further study.

We reconstituted *RIF1^-/-^* U-2 OS cells with full-length RIF1-L and RIF1-S coding sequences (CDS) cloned into a tetracycline-inducible pcDNA5-eGFP-FRT/TO plasmid vector (Addgene plasmid #19444) by Gateway recombination cloning (Invitrogen, 11789020 and 11791020) (6). Resulting GFP-RIF1-L and GFP-RIF1-S plasmids were cotransfected into *RIF1^-/-^*Tet-on U-2 OS cells with pOG44 using jetPRIME^®^, selected with 200 µg/ml hygromycin for one week, and tested for RIF1 expression following induction with 1 µg/ml doxycycline (Dox). GFP plasmid was included as a control. The three resulting cell lines, *RIF1^-/-^*:GFP, *RIF1^-/-^*:GFP-RIF1-L and *RIF1^-/-^*:GFP-RIF1-S are also being referred as GFP, GFP-RIF1-L and GFP-RIF1-S expressing cells throughout this paper. *RIF1^-/-^*U-2 OS cells expressing GFP-tagged RIF1^CTD^ constructs were generated by PCR amplifying codons 2170-2472 of the RIF1-L or RIF1-S CDSs followed by Gateway cloning into pcDNA5-eGFP-FRT/TO. Point mutations were introduced through QuikChange mutagenesis method with Phusion^®^ High Fidelity DNA Polymerase (Thermo Scientific, F530) or KOD Hot Start Polymerase (Novagen, 71086). All constructs were sequenced in their entirety. GFP-RIF1-L^CTD^ and GFP-RIF1-S^CTD^ constructs were stably transfected into *RIF1^-/-^* U-2 OS cells as described above. Experimentally verified CDK1 phosphorylation sites of human RIF1 protein were queried from UniProt website under the primary accession ID: Q5UIP0 on 6^th^ July 2022.

### Generation of RIF1-L-deficient mice

One-cell embryos from C57BL/6J mice were injected with a mixture of 40 ng/µl of Cas9 protein (PNA Bio) and 25 ng/µl of each of the two sgRNAs complementary to the 3’ portion of Rif1-In31 (sgRNA1: CCTAACATTTTACAAGGGCGATT) and 5’ portion of Rif1-Ex32 (sgRNA2: CCCAGGATCACAGAGCTCTAAAT) of the m*Rif1* gene spanning nucleotides 51962725 to 52016781 of mouse chromosome 2 (Reference GRCm39 C57BL/6J). The region flanked by sgRNA1 and sgRNA2 comprised of 149 nt that includes Rif1-Ex32 5’ splice acceptor (SA) site. Tail DNA from founder mice was subjected to deep sequencing using m*Rif1* primers and those mice exhibiting *Rif1* mutations were backcrossed to wild-type mice to obtain germline m*Rif1* mutant lines. Two lines were selected to be used in this study: a *Rif1^iA^* line harboring a single nucleotide (A) insertion at codon 2010 that likely reflects cleavage and error-prone repair at sgRNA2, and a *Rif1^ΔEx32^* line harboring a 129 nt deletion between sgRNA1 and sgRNA2 that deletes the 3’portion of intron 31 and the 5’ portion of Ex32, including the 5’ SA site. Analysis of T and B cell development, mitogenesis, and *in vitro* class switch recombination (CSR) were carried out using cells isolated from bone marrow and spleen of 6-8-week old *Rif1^+/+^*and *Rif1^ΔEx32/ΔEx32^* mice as previously described (86, 87).

### RNAi and shRNA screening

siRNA screen for RIF1 splicing regulators was performed through a SMARTpool siRNA library (Dharmacon) consisting of 151 siRNAs targeting RNA binding proteins in the human genome that was obtained through UW Small Molecule Screening & Synthesis Facility (SMSSF) (88). Approximately 10,000 HeLa cells were reverse-transfected with 20 nM of each of the siRNAs using DharmaFECT1 (Dharmacon) in 96-wells plates for a 48-hour period (Sup. Fig. 4A). Subsequently, the transfected cells were lysed in TRIzol reagent for RNA extraction, and cDNA was synthesized as described. RIF1 splicing assay was performed as described to identify candidate RIF1 splicing factors.

The following shRNA lentiviral vectors targeting candidate RIF1 splicing regulators were purchased from Sigma: SRSF1 (cat# TRCN0000001095); SRSF2 (cat# TRCN0000000084); SRSF3 (cat# TRCN0000001227); SRSF7 (cat# TRCN0000001142); PTBP1 (cat# TRCN0000231420); RBM28 (cat# TRCN0000239461); and snRNP70 (cat# TRCN0000000011).

Non-targeting vector (Addgene plasmid #1864) was included as negative control. Lentiviral particles were produced by transient transfection of HEK293T cells with shRNA vectors, psPAX2 (Addgene plasmid #12260) and pCMV-VSV-G (Addgene plasmid #8454) in a ratio of 3:2:1 by jetPRIME^®^ as described (83, 89). Viral supernatants harvested at 24 h and 48 h post-transfection were incubated with U-2 OS cells for 24 h followed by selection in media containing 2 µg/ml puromycin for 48-72 h. Cells were harvested in TRIzol reagent for RIF1 splicing assays.

### RNA immunoprecipitation (RNA-IP)

Approximately 50 million cells were lysed in 1 mL ice-cold NET-2 buffer (50 mM Tris-HCl (pH 7.5), 150 mM NaCl, 0.05% v/v NP40) supplemented with 2 mM 1,4-dithiothreitol, 0.2 U/µL RNasin Plus (Promega, N2611), 20 mM sodium fluoride, 20 mM β-glycerophosphate and 2X protease inhibitor cocktail (Sigma, P8340-5ml; Thermo Scientific, 78438). The lysate was sonicated using five pulses of 3 s ON, 30 s OFF followed by three pulses of 10 s ON and 30 s OFF at an amplitude of 30% (Fisher Scientific, FB120). The lysate was cleared by centrifugation at 14,000 *x* g for 10 min, 4°C. The supernatant was incubated with 5 µg of either the targeted antibodies (SRSF3: Cell Signaling Technology #35073; SRSF7: Bethyl A303-772A; PTBP1: ThermoFisher 32-4800; SRSF1-M: Santa Cruz sc33652; SRSF1-R: Abcam ab38014; SRSF2: Proteintech 20371-1-AP; snRNP70: Invitrogen #PA5-115943) or normal IgG controls (Mouse: Millipore 12-371; Rabbit: Millipore 12-370) for 1 h on a nutator mixer at 4°C before Protein A/G PLUS-Agarose bead suspension (Santa Cruz, sc-2003) was added (20 µl/1 µg of antibody) for overnight incubation. The beads were washed with NET-2 buffer five times and the immunoprecipitated RNA was extracted by TRIzol reagent. Relative fold enrichment of RIF1 pre-mRNA in the target RNA-IP sample versus the control IgG sample or the percent input was quantified by qPCR assay with two pairs of intron-exon primers. Primer set 1 (PS1) targets RIF1-In31 and RIF1-Ex32 while primer set 2 (PS2) targets RIF1-Ex32 and RIF1-In32 (RIF1-In31-F: 5’- TAGTCATCTAGGGTTCTGAGTG-3’ and RIF1-Ex32-R: 5’-TCCTTTGGCTGAAGTGGTATTATG- 3’; RIF1-Ex32-F: 5’-CATAATACCACTTCAGCCAAAGG-3’ and RIF1-In32-R: 5’- GTGACATGAAAACTAAAGCACTTC-3’).

### DNA replication pattern analysis

DNA replication pattern analyses were performed as described (83). *RIF1^-/-^*, *RIF1^-/-^*:GFP, *RIF1^-/-^*:GFP-RIF1-L, and *RIF1^-/-^*:GFP-RIF1-S U-2 OS cells were pulse labeled with 20 µM 5-ethynyl-2’- deoxyuridine (EdU) for 20 min and stained for EdU incorporation. The presence of early, mid, or late DNA replicative stages were accessed from the EdU staining patterns. The percentages were calculated for each sample. Alternatively, cells were synchronized with 2 mM thymidine for 19 h, released into thymidine-free growth media for 9 h, and then returned to thymidine-containing media for 6 h, at which time most cells are in mid-S phase (83). Cells were then pulse labeled with EdU as described (83). A minimum of 100 cells per sample was imaged by confocal microscopy.

### EdU labeling, flow cytometry, immunofluorescent microscopy and live cell imaging

For cell cycle progression experiments, U-2 OS cells were incubated with 20 µM EdU for 30 min before collection and then fixed with ice-cold 70% ethanol. EdU detection was performed using the Click-IT Plus EdU Alexa Fluor 647 Flow Cytometry Assay Kit (Life Technologies, C10634).

Propidium iodide (PI) was added at a concentration of 50 µg/ml. Flow cytometry was performed on Thermo Fisher Attune, and data was analyzed and organized using FlowJo software. For *in situ* EdU and 5-bromo-2′-deoxyuridine (BrdU) staining, U-2 OS cells were pulse labelled with 20 µM BrdU or EdU for 30 min and fixed with 4% w/v paraformaldehyde (PFA). For BrdU detection, cells were then incubated with 2 M HCl for 30 min and then permeabilized with 0.2% v/v Triton X- 100 for 15 min at room temperature, washed and blocked in 3% w/v bovine serum albumin (BSA). Cells were stained with BrdU primary antibody (Santa Cruz, sc-32323) in 3% BSA and incubated overnight at 4°C, followed by washing in 0.02% v/v PBST (PBS with 0.02% v/v Tween-20) and incubation with appropriate secondary antibodies in 3% BSA for 1 h at room temperature. EdU was detected by click chemistry as described above. Samples were mounted in VECTASHIELD mounting medium with DAPI (Vector, H-1200) before imaging.

For immunostaining experiments, cells were seeded into 12-well plate with glass coverslips, fixed with 4% w/v PFA, permeabilized with 0.2% v/v Triton X-100 and blocked with 3% w/v BSA at room temperature. The coverslips were then transferred to an improvised humidity chamber for immunostaining with the appropriate primary antibodies at 37°C for an hour or at 4°C overnight. Primary antibodies and the dilution factors used were listed in figure legends. Coverslips were washed 3 X 10 min with 0.05% v/v PBST (PBS with 0.05% v/v Tween-20) before incubating with Alexa Fluor™ secondary antibodies (Thermo Fisher #A11032, #A32733) at a dilution factor of 1:10,000 at room temperature for 45 min. Coverslips were washed 3 X 10 min with 0.05% PBST followed by 2 X 1 min with PBS before mounting. Nuclear DNA was either stained with 0.5 µg/ml DAPI for 10 min at room temperature and then mounted with mounting medium for fluorescence (Vector, H-1000), or directly mounted in mounting medium with DAPI for fluorescence (Vector, H- 1200) before imaging. Images were acquired using Nikon A1RS or Nikon AX confocal microscopes under the desired objectives with AU = 1. Images were organized using NIS-Elements Advanced Research/Fiji ImageJ software. Foci counts were done in CellProfiler (version 4.2.6) or ImageJ macro Foci_Analyzer_1_5 (https://imagej.net/plugins/foci-analyzer). Anisosome area measurement was performed in Fiji utilizing Labkit classifier followed by manual annotation. Outputs were extracted and visualized in R, MATLAB, and/or Prism.

For live cell imaging, cells were plated into 38 mm glass bottom dishes and stained with 1 µg/mL of Hoechst 33342 for 30 min before being imaged in humidified CO_2_ chamber supplemented with 5% CO_2_ throughout the experiment.

### *In vitro* DNA binding assay

Fluorescein-labelled antiparallel G4 DNA (5ʹ-FAM-TTT TTT GGG GGG GGG GGG GGG GGG GG-3ʹ) was folded in refolding buffer (1 µM DNA, 10 mM Tris-HCl (pH 7.5), 50 mM KCl, 1 mM ethylenediaminetetraacetic acid (EDTA), 40% v/v polyethylene glycol 200), by heating to 95°C for 10 minutes and subsequent cooling over 4 hours. Purified RIF1 CTD was incubated with 5 nM DNA in reaction buffer (20 mM HEPES (pH 7.6), 50 mM KCl, 1 mM EDTA, 0.01% v/v Triton X- 100, 10 % v/v polyethylene glycol 200) for 30 minutes at 30 °C. The fluorescence anisotropy of each sample was measured at 25°C with a Beacon 2000 fluorescence polarization system.

### Protein extraction, chromatin fractionation and immunoblotting

For whole-cell protein extraction, cells were lysed in either high salt lysis buffer (50 mM Tris (pH 7.5), 300 mM NaCl, 10% glycerol, 2 mM MgCl_2_, 3 mM EDTA, and 0.5% Triton X-100) or modified RIPA buffer (50 mM Tris-HCl (pH 7.4), 150 mM NaCl, 1 mM EDTA, pH 8, 1% sodium deoxycholate, 0.1% SDS, and 1% Triton X-100) supplemented with 10 mM sodium fluoride, 10 mM β-glycerophosphate and 1X Protease Inhibitor Cocktail (Sigma, P8340-5ml; Thermo Scientific, 78438). The lysate was incubated on ice for 20 min and sonicated using 5 pulses of 3 s ON, 5 s OFF at an amplitude of 30% (Fisher Scientific, FB120) before the addition of 4X SDS sample buffer (200 mM Tris-HCl (pH 6.8), 40% glycerol, 8% SDS, 0.5% bromophenol blue and 10% beta-mercaptoethanol). The samples were heated at 95°C for 5 min prior to freezing at -20°C for storage or loading directly for immunoblotting.

For chromatin fractionation of full-length RIF1, cells were resuspended in CSK buffer (20 mM HEPES (pH 7.4), 150 mM NaCl, 3 mM MgCl_2_, 300 mM sucrose, and 0.5% Triton X-100) supplemented with 20 mM sodium fluoride, 20 mM β-glycerophosphate and 2X Protease Inhibitor Cocktail. The cells were incubated on ice for 20 min. Fifty percent of the cell suspension was kept as whole cell extract (WCE) with the addition of 50 U/ml Benzonase (Sigma, E1014-5KU) followed by a 20 min on-ice digestion. The remaining cell suspension was centrifuged for 5 min at 5,000 x *g* at 4°C. The supernatant was transferred to a new tube and saved as the soluble fraction (SF) which contains cytoplasmic and nucleoplasmic content, while the pellet/chromatin fraction (CF) was washed twice in CSK buffer without detergent and resuspended in complete CSK buffer with 50 U/ml Benzonase for a 20 min on-ice digestion. All lysates were mixed with 0.5 v/v of 4X SDS sample buffer and heated at 95°C for 15 min before loading. Chromatin fractionation of the RIF1 CTD constructs was done with the same procedure, but the salt concentration of the CSK buffer was reduced to 100 mM due to the higher solubility of RIF1 CTD compared to full-length RIF1.

For immunoblotting, samples were separated by 6%, 12% or 15% SDS-polyacrylamide gel (SDS-PAGE) depending on the molecular weight of the target protein and transferred to 0.45 µm Immobilon®-FL PVDF membranes (MilliporeSigma, IPFL00010). For small molecular weight RBP transfer, 0.2 µm Amersham™ Protran® Western blotting nitrocellulose membranes (MilliporeSigma, GE10600001) was used in Tris-glycine transfer buffer supplemented with 10% v/v methanol. For large molecular weight transfer, 0.01% v/v SDS was added in the transfer buffer. The membranes were blocked with blocking solution (5% w/v milk in Tris-buffered saline, 0.1% v/v Tween 20 (TBST)) for an hour before blotting with target primary antibodies overnight at 4°C. The source and the dilution of the primary antibodies used were listed as followed: Total RIF1 (Bethyl Laboratories A300-569A; 1:500); GFP (Santa Cruz sc9996, 1:100); MCM2 (Santa Cruz sc373702, 1:100 or Abcam ab4461, 1:1000); MCM4 (Abcam ab4459, 1:1000); vinculin (Santa Cruz sc73614, 1:100); SRSF1 (ab38014, 1:1000); SRSF3 (Cell Signaling Technology #35073, 1:1000); SRSF7 (Bethyl A303-772A, 1:1000); PTBP1 (ThermoFisher 32-4800, 1:1000); GAPDH (Cell Signaling Technology #2118S); lamin B1 (Abcam ab16048, 1:2000); α-tubulin (Sigma T6199, 1:500); and β-tubulin (Sigma Millipore 05-661, 1:1000). After that, the membranes were washed 3 X 5 min with TBST and incubated with LI-COR IRDye secondary antibodies (IRDye 800CW goat anti-rabbit or IRDye 680RD goat anti-mouse) at a dilution of 1:10000 in blocking solution for an hour at room temperature. Membranes were washed 3 X 5 min with TBST and images were acquired using Odyssey Fc/XF (LI-COR Biosciences). The exported images were then analyzed and organized with ImageStudio software (v5.2, LI-COR Biosciences).

### RIF1 isoform-specific antibodies

α-RIF1-L, α-RIF1-S, and α-RIF1-pS2260/65 antibodies (Lifetein, LLC) were peptide-affinity purified from the sera of rabbits injected with the following KLH-conjugated immunogens, respectively: hRIF1-S (N-VKTSPTTQSKISEMAKESIP-C); hRIF1-L (N- AKGFLSPGSRSPKFKSSKKC-C); RIF1-pS2260/65 (N-AKGFL[pS]PGSRPKFKSSKKC-C). α-RIF1-pS2260/65 antisera were first immunodepleted against an identical non-phosphorylated peptide prior to affinity purification using the phosphopeptide. All antibodies were used at a dilution of 1:500 for Western blotting and immunofluorescence staining experiments.

### RIF1 purification and mass spectrometry (MS)

RIME (rapid immunoprecipitation mass spectrometry of endogenous proteins) assay of GFP- RIF1-L and GFP-RIF1-S expressed on a *RIF1^-/-^* background was carried out as described (41, 83) (Fig. 5A). Briefly, ∼20 million cells were counted and fixed with 20 ml 1% formaldehyde solution for 8 minutes at room temperature. Fixation was quenched by adding 0.12 M glycine. The soluble fraction was extracted in 10 ml of LB1 (50 mM HEPES-KOH (pH 7.5), 140 mM NaCl, 1 mM EDTA, 10% Glycerol, 0.5% NP-40, 0.25% Triton X-100, 1X Protease Inhibitor Cocktail (Sigma, P8340- 5ml) for 10 min with rotation at 4°C. Cell nuclei were pelleted and washed once with 10 ml LB2 (10 mM Tris-HCl (pH 8.0), 100 mM NaCl, 1 mM EDTA, 0.5 mM EGTA, 1X Protease Inhibitor Cocktail) and then resuspended in 500 µl LB3 (10 mM Tris-HCl (pH 8.0), 100 mM NaCl, 2.5 mM MgCl_2_, 0.1% w/v sodium deoxycholate, 0.5% Triton X-100, 1X Protease Inhibitor Cocktail) with 500 U Benzonase and incubated at room temperature for 30 min. Benzonase was deactivated with 2 mM EDTA and 1 mM EGTA. The mixture was supplemented with 50 µl 10% Triton X-100 and 37.5 µl of 4 M NaCl before LB3 was added to bring the total lysate volume of each sample to 1 ml. Digested lysates were sonicated for 3 pulses of 10 s ON, 50 s OFF at an amplitude of 40% and clarified by centrifugation at 20,000 x *g* for 10 min at 4°C. The supernatants were incubated with ChromoTek GFP-Trap Magnetic Agarose beads (Fisher Scientific) per manufacturer’s recommendations on a nutator mixer at 4°C overnight. Subsequently, 50 µl of pre-washed Dynabeads protein G (Invitrogen, 10003D) was added to the lysates and incubated for additional 4 h at 4°C.

For Western blotting, beads were washed sequentially with 1 ml LB3 and 1 ml RIPA buffer (50 mM HEPES-KOH (pH 7.5), 0.5 M LiCl, 1 mM EDTA, 1% NP-40, 0.7% w/v sodium deoxycholate, 1X Protease Inhibitor Cocktail) and boiled in 100 µl 2X SDS sample buffer. For mass spectrometry, beads were washed 5 times with 1 ml of RIPA buffer and twice in 1 ml of cold freshly prepared 100 mM ammonium hydrogen carbonate (AMBIC) solution and processed as described (41).

GFP-RIF1 RIME IPs were subjected to tryptic digestion and LC-MS/MS analysis using an Orbitrap Fusion™ Lumos™ Tribrid™ mass spectrometer using the filter aided sample preparation (FASP) method (90). The tryptic digest solution was desalted/concentrated using an Omix 100 µL (80 µg capacity) C18 tip. The solution was pipetted over the C18 bed 5 times, and rinsed 3 times with H_2_O, 0.1% trifluoroacetic acid (TFA) to desalt. The peptides were eluted from the C18 resin into 150 µL 70% acetonitrile, 0.1% TFA and lyophilized. The peptides were re-suspended in 95:5 H_2_O:acetonitrile, 0.2% formic acid and analyzed in duplicate as described below. Samples were analyzed using a UPLC-MS/MS system consisting of an Easy-nLC 1200 ultra-high-pressure liquid chromatography system and an Orbitrap Fusion Lumos mass spectrometer (ThermoFisher Scientific). Peptides were loaded in buffer A (H_2_O, 0.2% formic acid) at a pressure of 300 Bar onto a 20-cm-long fused silica capillary nano-column packed with C18 beads (1.7 μm-diameter, 130 Angstrom pore size from Waters). Peptides eluted over 120 minutes at a flow rate of 350 nL/min with the following gradient established by buffer A (H_2_O, 0.2% formic acid) and buffer B (80% acetonitrile (ACN), 0.2% formic acid): Time/T = 1 mins, 5% buffer B; T = 52 mins, 30% buffer B; T = 80 mins, 42% buffer B; T = 90 mins, 55% ACN; T = 95 to 100 mins, 85% buffer B; T = 101 to 120 mins, equilibrate at 0% buffer B. The nano-column was held at 60°C using a column heater (in-house constructed) (91).

The nanospray source voltage was set to 2200 V. Full-mass profile scans were performed in the orbitrap between 375-1500 m/z at a resolution of 120,000, followed by MS/MS HCD scans in the orbitrap of the highest intensity parent ions in a 3 seconds cycle time at 30% relative collision energy and 15,000 resolution, with a 2.5 m/z isolation window. Charge states 2-6 were included and dynamic exclusion was enabled with a repeat count of one over a duration of 30 seconds and a 10 ppm exclusion width both low and high. The AGC target was set to “standard”, maximum inject time was set to “auto”, and 1 µscan was collected for the MS/MS orbitrap HCD scans.

The MetaMorpheus software program was used to identify peptides and proteins in the samples (92, 93). Protein fold changes were normalized and quantified from two technical replicates for each of the three biological replicates by FlashLFQ (94, 95).

### Proximity-ligation assay (PLA)

*RIF1^-/-^* U-2 OS cells expressing GFP, GFP-RIF1-L, or GFP-RIF1-S were mock irradiated or exposed to 10 Gy IR followed by 2 h recovery. The cells were then pre-extracted for 8 min with CSK buffer containing 100 mM NaCl and 0.5% v/v Triton X-100 to reduce cytoplasmic background signal prior to cell fixation and primary antibodies incubation (α-GFP, Santa Cruz sc9996, 1:250; α-MDC1, Sigma HPA006915, 1:500) overnight. Reagent kits for Duolink® Proximity Ligation Assay (Sigma) were used, and PLA was performed according to the manufacturers’ conditions. PLA foci number for each genotype was quantified by customized R scripts.

### Neutral Comet Assay

*RIF1^-/-^* U-2 OS cells expressing GFP, GFP-RIF1-L, or GFP-RIF1-S were mock irradiated or exposed to 10 Gy IR and harvested immediately (T0) or allowed for an hour of recovery in 37°C (T1). Neutral Comet Assay was performed according to Trevigen’s protocol. Cells were harvested in 1X PBS (Trevigen #4870-500) and combined with molten CometAssay LMAgarose (Trevigen #4250-050-02) at 37 °C at a ratio of 1:10 (v/v) at a cell density of ∼1 x 10^5^/mL. For each sample, 50 µL of the cell suspension was added onto a well of the CometSlide™ (Trevigen #4250-050- 03). The slides were allowed for gelling for 30 min in the dark at 4°C before being incubated in Lysis Solution (Trevigen #4250-050-01) at 4°C overnight. After the slides were equilibrated in 1X TBE buffer (90 mM Tris, 90 mM boric acid, 2 mM Na_2_EDTA, pH 8.33) for 15 minutes, the nucleoids were subjected to electrophoresis at 22 V, 30 minutes at 4°C in the dark. The slides were then immersed in distilled water for 5 min followed by 5 min in absolute ethanol at room temperature. The slides were subsequently dried at 37 °C for 15 min and then kept at room temperature for storing. Prior to imaging, the nucleoids were stained with 10 µg/mL propidium iodide for 30 min at room temperature. The slides were immersed in distilled water briefly, dried on paper towel, and imaged immediately. Comet images were segmented by OpenComet (96) in Fiji using the default parameter. Inaccurate segmentations were removed manually. Tail DNA percent of each segmented comet was calculated by OpenComet and extracted from its output using a custom MATLAB script.

### *In vitro* phase separation assay

RIF1-L and RIF1-S CTD fragments were subcloned into pDEST15 for expression as glutathione-S-transferase (GST) fusion proteins. GST-tagged RIF1 CTD fragments were transformed into BL21-AI strain. Protein expression was induced by growing the transformed bacteria in TB medium supplemented with 1 mM IPTG + 0.2% w/v arabinose and grown at 16°C overnight. Cells were then lysed in lysis buffer (50 mM HEPES (pH 7.5), 300 mM NaCl, 1 mM DTT, 100 mM dextrose, 10% glycerol, 1X Protease Inhibitor Cocktail) by sonication. The soluble fractions were collected by centrifugation at 20,000 x g for 30 minutes at 4°C. The proteins were purified with GS4B resin and eluted in GS4B elution buffer (20 mM HEPES (pH 7.4), 300 mM NaCl, 10% glycerol, 20 mM reduced glutathione, 200 mM trehalose). Purified GST-RIF1^CTD^-L and GST- RIF1^CTD^-S were then subjected to phase separation assays in the presence of 10% dextran/polyethylene glycol (PEG) as described (97). Images were collected at 10 min post-mixing.

### Fluorescence recovery after photobleaching (FRAP)

U-2 OS cells were seeded in 38 mm glass bottom dishes and transfected at 60-80% confluency with 2.5 µg of GFP-RIF1^CTD^ plasmid using Lipofectamine 3000 (ThermoFisher, L3000015) in 7.5 µL Lipofectamine and 5 µL P3000 reagents, each diluted in 120 µL serum-free McCoy’s 5A medium. The transfection medium was changed at 4 h post-transfection and the cells were incubated overnight for transgene expression. Cells were kept in humidified CO_2_ chamber supplemented with 5% CO_2_ throughout the experiment. FRAP was programmed through ND Stimulation module (Nikon NIS-Elements AR) on Nikon AX confocal microscope 24 hours post-transfection with 25% 488 nm laser for 1.02 s on anisosomes of roughly 2.5 nm in diameter. Recovery images were acquired at 5 s interval for 3 min post-bleaching. The resulting videos were analyzed in MATLAB scripts (https://github.com/adenine-koo/FRAP.git) according to the easyFRAP pipeline (98). Briefly, the average pixel values of three regions of interest (nucleus, bleached anisosome, and background) were extracted. Frame stabilization of the bleached anisosome was performed based on a correlation algorithm. Full scale normalization was done to correct background fluctuation, starting intensity difference, acquisition bleaching/laser fluctuation as well as variation in bleaching depth. Each recovery curve was plotted and fitted by a single exponential equation to estimate its t-half value and mobile fraction.

### Statistical processing

Statistical analysis information including individual biological and technical replicates number, mean or median, and error bars are explained in the figure legends. Statistical tests were performed in Prism (v10, GraphPad) or R. The tests performed and the resulting *p* values are listed in the figure legends.

## DATA AND MATERIALS AVAILABILITY

Raw data will be available upon request.

## SUPPORTING INFORMATION

This article contains supporting information.

## ACKNOWLEDGMENTS

The authors would like to thank Lance A. Rodenkirch (University of Wisconsin Optical Image Facility) for assistance with FRAP experiments and anything microscopy. We appreciate the help from Joan Steitz and Vanessa Mondol (Yale School of Medicine) in optimizing our RNA IP protocol. ASHK thanks Adrian Tan (National Parks Board Singapore) for his biostatistical inputs, and all the contributing users on image.sc forum including @scouser27 and @bramvdbroek for their valuable feedback in establishing the robust image analysis pipelines.

## AUTHOR CONTRIBUTIONS

Conceptualization, RST, ASHK, WYJ, MS, LMS, JLK, LG, CJB; Methodology, all; Investigation, Validation, and Formal Analysis, ASHK, WYJ, MS, CEB, YC, DW, AFV, AB; Software, Data Curation, and Visualization, ASHK, WYJ, MS, CEB, YC, DW, AFV, AB; Writing – Original Draft, RST, ASHK and WYJ; Writing – Review & Editing, all; Supervision, Resources, and Funding Acquisition, RST, LMS, DW, JLK, LG, CJB;

---

**Sup. Fig. 1.** Effects of canonical DNA repair inhibitors on CLM-dependent RIF1 AS. (A, left panel) U-2 OS cells treated with CLM with or without PARP inhibitor for 4 h were analyzed by RIF1 splicing assay in three biological replicates. (A, right panel) Quantification of the mean RIF1- L/S mRNA ratio ± standard error by densitometry from the left panel. Each dot represents an individual biological replicate, N = 3. (B, left panel) U-2 OS cells treated with CLM, ATM or DNA-PK inhibitor for 4 h were analyzed by RIF1 splicing assay in four biological replicates. (B, right panel) Quantification of the mean RIF1-L/S mRNA ratio ± standard error by densitometry. Each dot represents an individual biological replicate, N = 4.

**Sup. Fig. 2.** Comparison of T and B cell development in *Rif1^+/+^* and *Rif1^ΔEx32^* mice. (A) B cell development in the bone marrow (BM) of *Rif1^+/+^* and *Rif1^ΔEx32^* mice. BM cells from the mice were stained with anti-B220 and anti-IgM, anti-CD43 or anti-CD25 antibodies. The percentages of B220^+^IgM^-^ pro/pre-, B220^+^IgM^+^ immature and B220^hi^IgM^+^ mature B cells (Left), B220^+^CD43^+^ pro- B cells (Middle), and B220^+^CD25^+^ pre-B cells (Right) in the gated live cells were shown. (B) B cell maturation in the spleen of *Rif1^+/+^*and *Rif1^ΔEx32^* mice. Spleen cells from the mice were stained with anti-B220, anti-IgM, and anti-IgD or anti-B220, anti-CD21, and anti-CD23 antibodies. The percentages of B220^+^ cells in the gated live cells (Upper); transitional 1 (T1) (IgM^hi^IgD^lo^), transitional 2 (T2) (IgM^hi^IgD^hi^), follicular (FO) mature (IgM^lo^IgD^hi^) B cells (Middle) and marginal zone (MZ) (CD21^hi^CD23^lo^) B cells within gated B cell populations (Lower) were indicated. (C) T cell development in the thymus of *Rif1^+/+^*and *Rif1^ΔEx32^* mice. Thymocytes from the mice were stained with anti-CD4 and anti-CD8. The percentages of DN, DP, CD4, and CD8 T cells in the gated live cells were shown. (D) T cell subpopulations in the spleen of *Rif1^+/+^* and *Rif1^ΔEx32^* mice. Splenocytes from the mice were stained with anti-CD4, anti-CD8, anti-CD62L and CD44. The percentages of CD4 and CD8 T cells in the gated live cells (Upper) and CD62L^hi^CD44^lo^ naïve, CD62L^lo^CD44^hi^ effect memory, and CD62L^hi^CD44^hi^ central memory T cells in the gated CD4^+^ or CD8^+^ cells (Lower) were shown. (E) TCR-induced thymidine incorporation in *Rif1^+/+^* and *Rif1^ΔEx32^* T cells. Splenic CD4 and CD8 T cells sorted from the mice were stimulated with medium (med), anti-CD3, anti-CD3 plus IL-2, anti-CD3 plus anti-CD28, or PMA plus Ionomycin. Proliferative responses were determined by [^3^H]thymidine incorporation. (F) BCR-induced thymidine incorporation in *Rif1^+/+^* and *Rif1^ΔEx32^* B cells. Splenic B cells sorted from the mice were stimulated with medium (med), anti-IgM, or anti-IgM plus IL-4, anti-CD40, LPS, or PMA plus Ionomycin. Proliferative responses were determined by [^3^H]thymidine incorporation. The data were obtained from 5 *Rif1^+/+^* and 4 *Rif1^ΔEx32^* mice.

**Sup. Fig. 3.** Comparison of IgG class switch recombination (CSR) potential of *Rif1^+/+^* and *Rif1^ΔEx32^* mice. (A-B) Splenic B cells from *Rif1^+/+^* and *Rif1^ΔEx32^* mice stimulated for 4 days with CD40 plus IL-4 or LPS plus IL-4 were analyzed for *in vitro* CSR by FACS for IgG1. Percentages of cells in the gated B220^+^ population were shown. (B) Graphical representation of the percentages of IgG1^+^ cells from (A) with mean ± standard deviation. The data were obtained from 2 or 3 *Rif1^+/+^* and 2 *Rif1^ΔEx32^* mice. Each mouse was analyzed in duplicates, and each dot represents one replicate. (C-D) Splenic B cells from *Rif1^+/+^* and *Rif1^ΔEx32^* mice stimulated for 4 days with LPS were analyzed for *in vitro* CSR by FACS for IgG2b and IgG3. Percentages of cells in the gated IgG2b^+^ or IgG3^+^ population were shown. (D) Graphical representation of the percentages of IgG2b^+^ or IgG3^+^ cells from (C) with mean ± standard deviation. Each experiment includes 3 *Rif1^+/+^* and 2 *Rif1^ΔEx32^* mice. Each mouse was analyzed in duplicates, and each dot represents one replicate.

**Sup. Fig. 4.** **RNAi screen for RIF1 splicing regulators**. (A) Schematic of RNAi screen in HeLa cells with 151 siRNAs targeting genes for RNA binding proteins and a non-targeting (NT) siRNA control by reverse transfection. (B) RIF1-L/S mRNA ratio from HeLa cells transfected with the indicated siRNAs. Green labels denote candidate positive regulators of Ex32 inclusion; red labels denote putative inhibitors of Ex32 inclusion (see Sup. Table 1 for the numbering key used for sample ID). (C) Candidate RIF1 splicing regulators chosen for secondary screening by shRNA knockdown based on the changes in RIF1-L/S mRNA ratio compared to NT.

**Sup. Fig. 5.** Subcellular colocalization of RIF1 splicing regulators in response to DNA damage and expression level in primary cancers. (A) Representative images of HeLa cells treated with DMSO, 5 nM or 10 nM of CLM for 4 h before cell fixation and immunofluorescence staining with α-SRSF1 (Abcam ab38017, 1:250), α-SRSF3 (Santa Cruz sc13510, 1:100), α- SRSF7 (Bethyl Laboratories A303-772A, 1:1000), and α-PTBP1 (Invitrogen 32-4800, 1:250). Scale bar = 25 µm. (B) Median expression level in transcript per million (TPM) for RIF1 splicing regulators – SRSF1, SRSF2, SRSF3, SRSF7, and PTBP1 in four TCGA cancer types of interest (see Fig. 2 for detailed information of each dataset). Graphs replotted from expression data assessed through GEPIA (99).

**Sup. Fig. 6.** **RIF1-L and RIF1-S isoforms have similar activity in canonical measures of RIF1 function**. (A) Western blot analysis of *RIF1^+/+^* U-2 OS cells and *RIF1^-/-^*U-2 OS cells stably transfected with plasmid vectors encoding GFP, GFP-RIF1-L, or GFP-RIF1-S. Both α-GFP (Santa Cruz sc9996, 1:200) and α-RIF1 (Bethyl Laboratories A300-569A; 1:500) detected a band of ∼300 kDa from *RIF1^-/-^*:GFP-RIF1-S and *RIF1^-/-^*:GFP-RIF1-L U-2 OS cells. Lamin B1 (Abcam ab16048, 1:2000) was included as loading control. (B) RIF1-L and RIF1-S are recruited to IR-induced foci with comparable efficiency. *RIF1^-/-^*:GFP-RIF1-S and *RIF1^-/-^*:GFP-RIF1-L U-2 OS cells were exposed to 10 Gy IR and stained with DAPI three hours post-irradiation prior to imaging. (C) MCM4 (Santa Cruz sc28317, 1:100 dilution) hyperphosphorylation in *RIF1^-/-^*U-2 OS cells were rescued by stable expression of either GFP-RIF1-S or GFP-RIF1-L. (D, E) RIF1-L and RIF1-S rescued the DNA replication pattern defect of *RIF1^-/-^* U-2 OS cells. Asynchronous U-2 OS cells of the indicated genotypes in (E) were pulse-labeled with EdU for 20 min and scored for the presence of early, mid, or late EdU staining patterns, as depicted in panel (D). (E) Representative EdU incorporation patterns of *RIF1^-/-^*:GFP-RIF1-S and *RIF1^-/-^*:GFP-RIF1-L U-2 OS cells. Cells exhibiting early and mid S-phase EdU incorporation patterns are denoted by green and yellow arrows respectively. (F) Quantification of the percentage of cells in early, mid, and late S-phase patterns as shown in (D) using a minimum of 100 cells per genotype. Note the lack of mid S-phase replication patterns in *RIF1^-/-^*cells that were rescued by both RIF1-L and RIF1-S. (G) RIF1^CTD^-S and RIF1^CTD^-L binds to antiparallel G4 quadruplex substrate with equal anisotropy *in vitro*.

## SUPPLEMENTARY TABLES

https://github.com/adenine-koo/RIF1_Raw-Sup_data

**Sup.** **Table 1**. siRNA key for the numbering used in Sup. Fig. 4B.

**Sup.** **Table 2**. Sheet 1 showed the results of student t-test performed on each of the 2784 genes between α-GFP-RIF1-L and α-GFP-RIF1-S immunoprecipitates (IPs) with the 94 significantly enriched genes selected and grouped in Sheet 2. Sheets 3 and 4 showed the student t-test results for the 2784 genes for α-GFP-RIF1-L or α-GFP-RIF1-S with α-GFP IP. Sheet 5 showed the results of two-way ANOVA with Tukey’s multiple comparisons that were performed on the 378 significantly enriched genes between α-GFP-RIF1-L, α-GFP-RIF1-S and α-GFP IPs.

## SUPPLEMENTARY VIDEOS

https://github.com/adenine-koo/RIF1_Raw-Sup_data

**Sup. Video 1.** Three-dimensional reconstruction of RIF1^CTD^-S anisosomes which resembled oblong spheroids from 10x 1 µm image sections taken at ∼ 25 s interval for a total of 534 s.

**Sup. Video 2.** Three-dimensional reconstruction of RIF1^CTD^-S anisosomes from 7x 2 µm image sections taken at 10.7 s interval for a total of 311 s, showing the dynamics of anisosomes and fusion events.

**Sup. Video 3.** RIF1^CTD^-L^5KQ^ (and other RIF1^CTD^-L variants) formed nested anisosome structures in which at least one smaller anisosome was formed within a bigger anisosome. Acquisition time interval = 1.1 s, scale bar = 10 µm. Note that this is a video from FRAP experiment so there was a bleaching event of 2 s before Frame #3.

**Sup. Video 4.** FRAP video images of RIF1^CTD^-S. Red arrow marked the bleached anisosome. Scale bar = 10 µm.

**Sup. Video 5.** FRAP video images of RIF1^CTD^-L. Red arrow marked the bleached anisosome. Scale bar = 10 µm.

