## Supplementary Figures for "DNA-damage dependent isoform switching modulates RIF1 DNA repair complex assembly and phase separation"

### Supplemental Figures

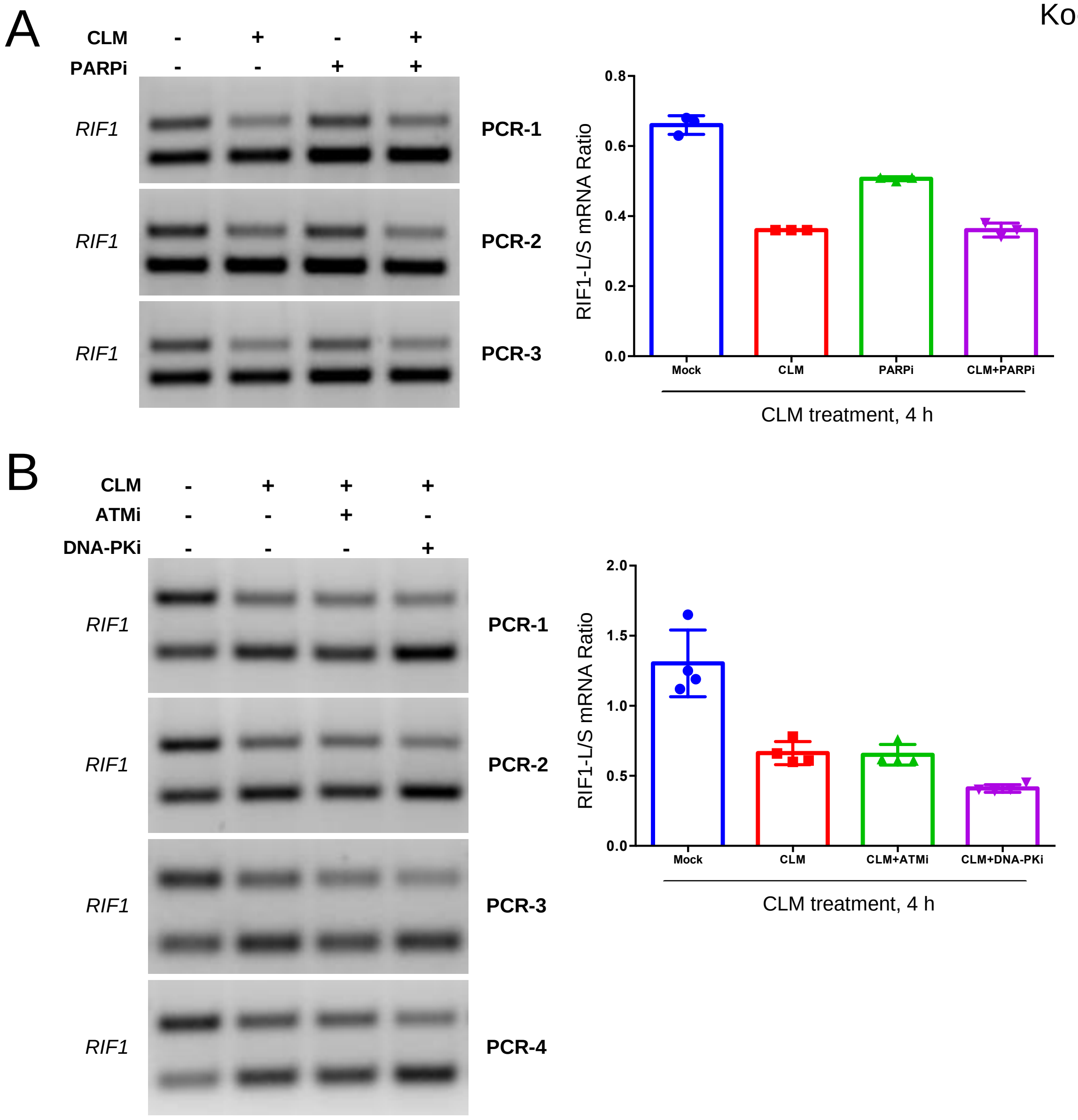
A
CLM - + - +
PARPi - - + +
RIF1
PCR-1
RIF1
PCR-2
RIF1
PCR-3
RIF1-L/S mRNA Ratio
0.0
0.2
0.4
0.6
0.8
Mock
CLM
PARPi
CLM+PARPi
CLM treatment, 4 h
B
CLM - + + +
ATMi - - + -
DNA-PKi - - - +
RIF1
PCR-1
RIF1
PCR-2
RIF1
PCR-3
RIF1
PCR-4
RIF1-L/S mRNA Ratio
0.0
0.5
1.0
1.5
2.0
Mock
CLM
CLM+ATMi
CLM+DNA-PKi
CLM treatment, 4 h

Koo *et al.* Sup. Fig. 2

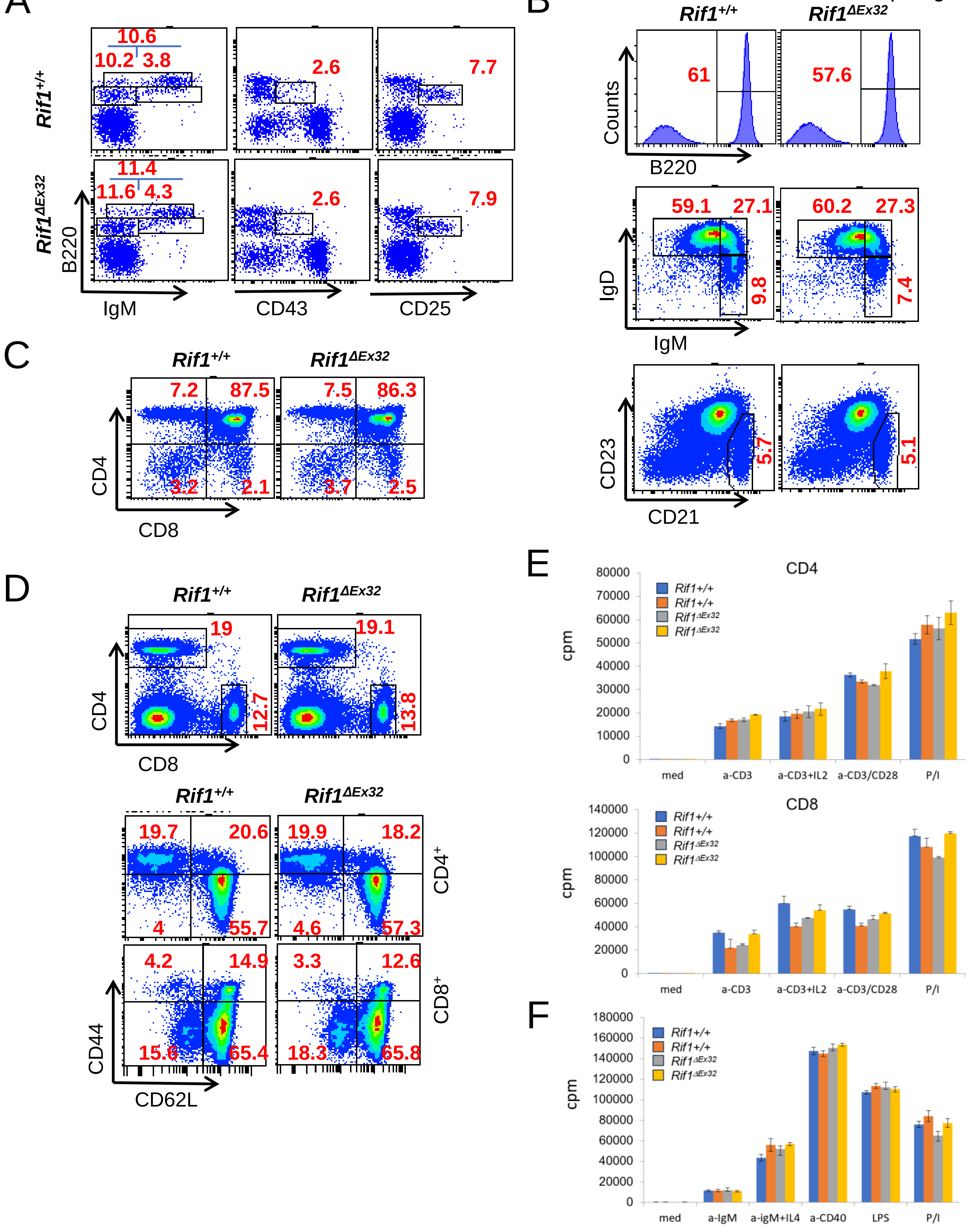

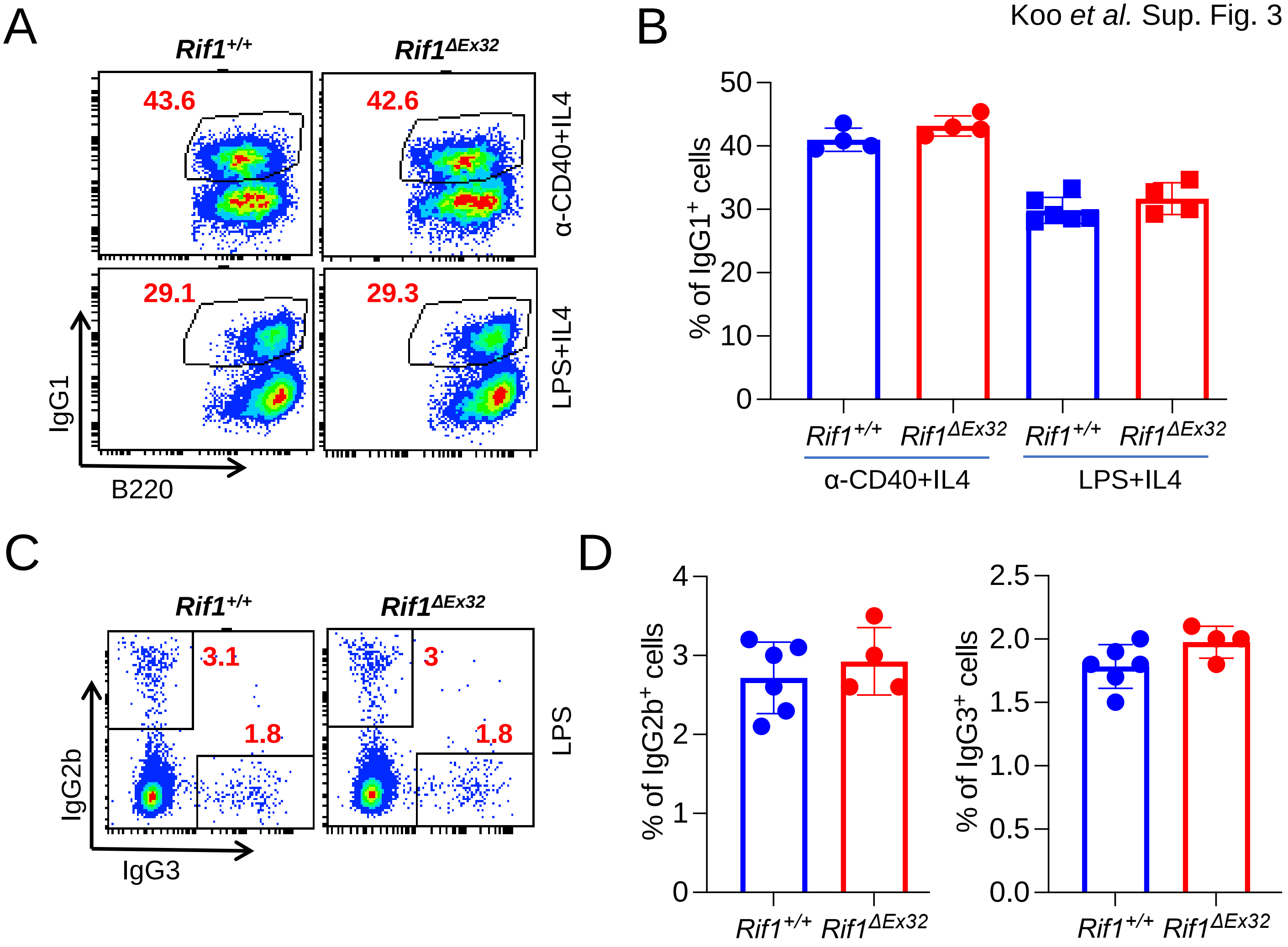
A
Rif1+/+
Rif1ΔEx32
43.6
42.6
29.1
29.3
α-CD40+IL4
LPS+IL4
IgG1
B220
B
50
40
30
20
10
0
% of IgG1+ cells
Rif1+/+
Rif1ΔEx32
Rif1+/+
Rif1ΔEx32
α-CD40+IL4
LPS+IL4
C
Rif1+/+
Rif1ΔEx32
3.1
1.8
3
1.8
LPS
IgG2b
IgG3
D
4
3
2
1
0
% of IgG2b+ cells
Rif1+/+
Rif1ΔEx32
2.5
2.0
1.5
1.0
0.5
0.0
% of IgG3+ cells
Rif1+/+
Rif1ΔEx32

A

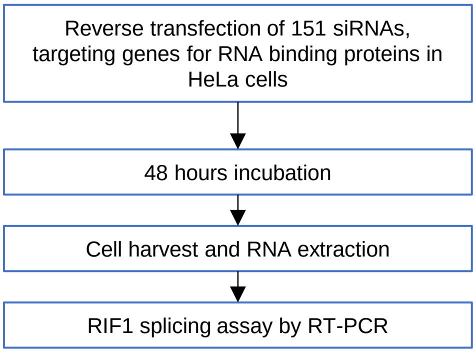


B

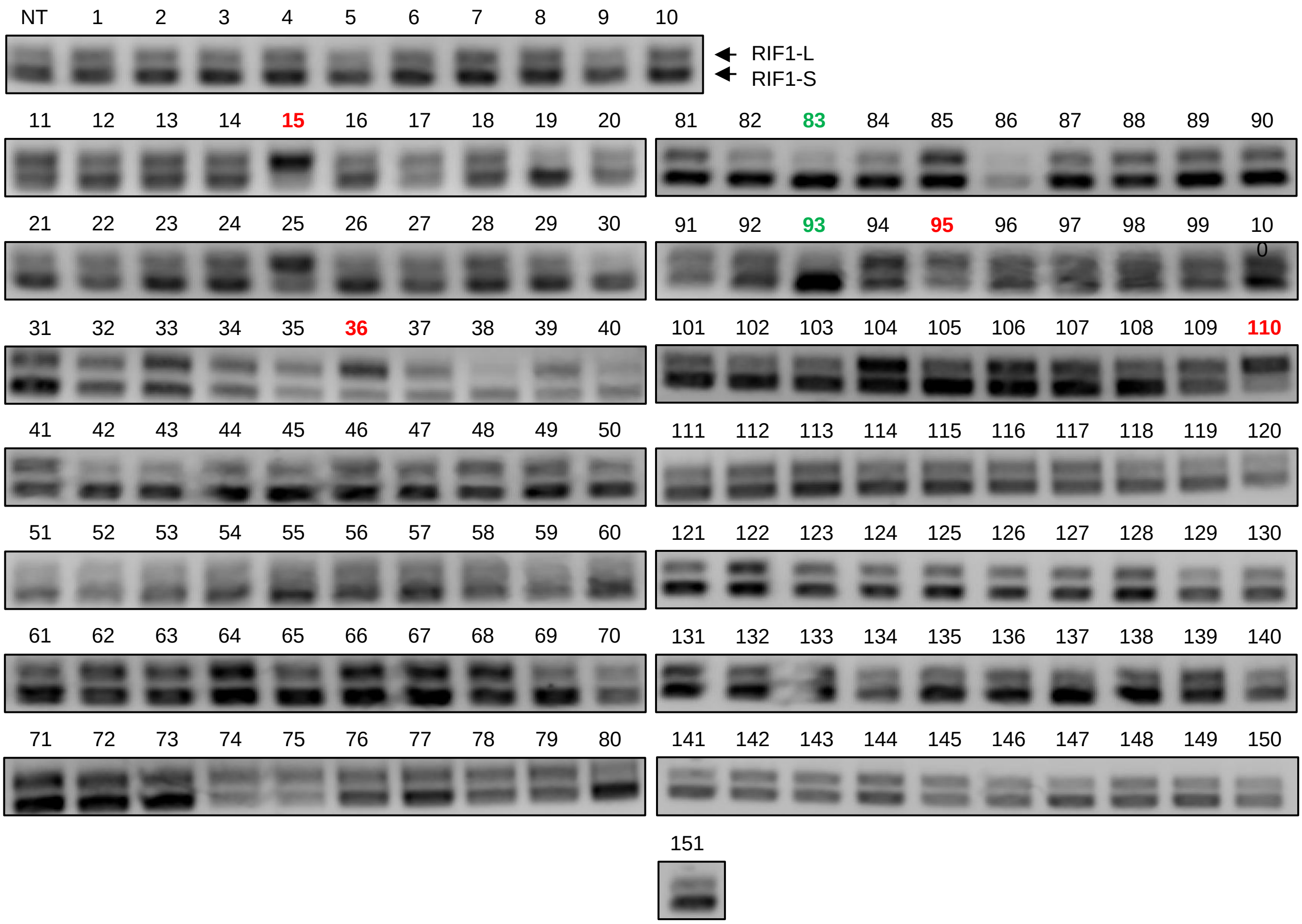


C

| Sample ID | Target | RIF1-L/S ratio |
|---|---|---|
| NT | Non-targeting control | 0.54 |
| 15 | PTBP1 | 2.61 |
| 36 | RBM28 | 1.33 |
| 83 | SRSF1 | 0.16 |
| 93 | snRNP70 | 0.25 |
| 95 | SRSF7 | 1.5 |
| 110 | SRSF3 | 2.83 |

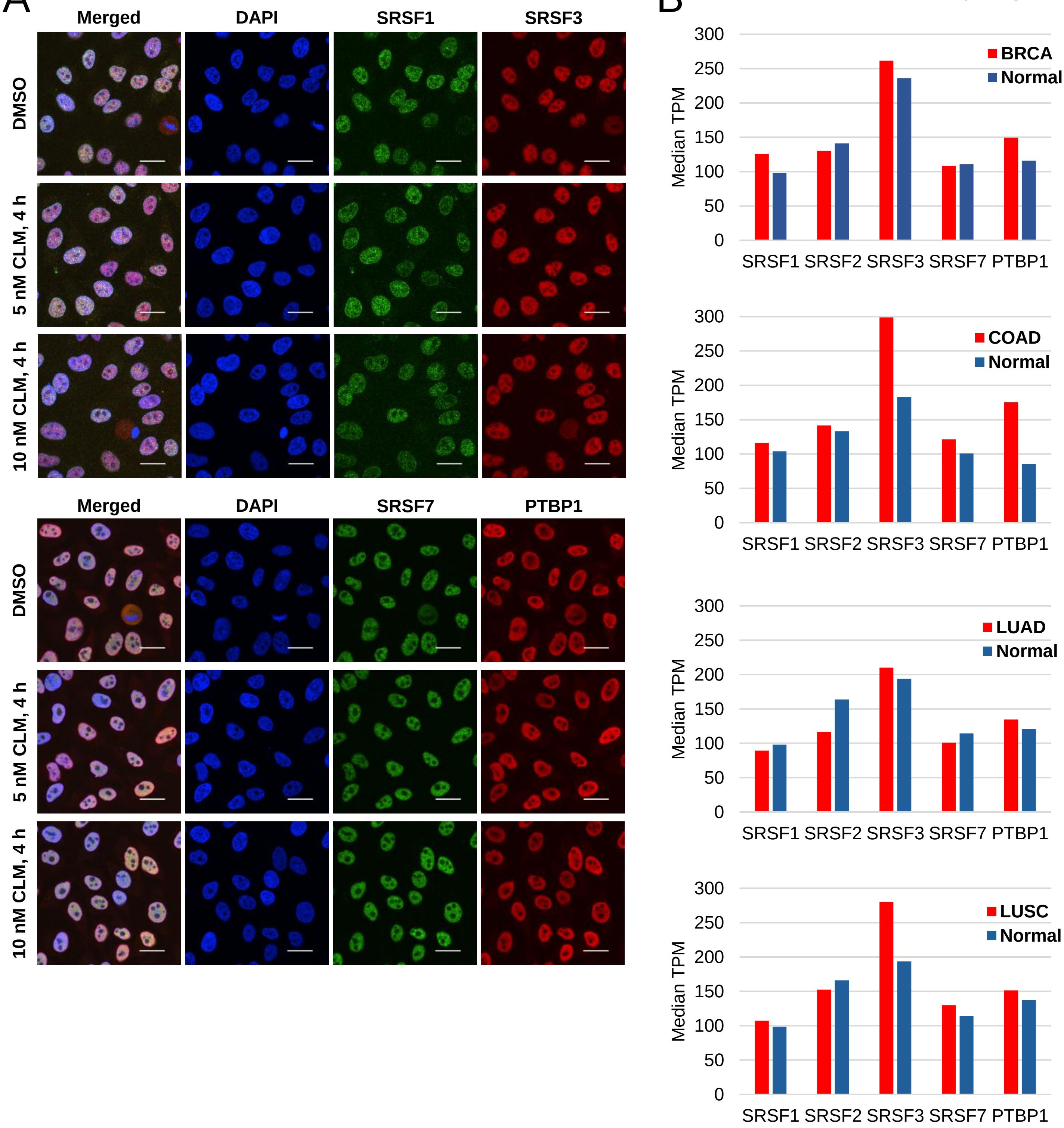
A
Merged
DAPI
SRSF1
SRSF3
DMSO
5 nM CLM, 4 h
10 nM CLM, 4 h
Merged
DAPI
SRSF7
PTBP1
DMSO
5 nM CLM, 4 h
10 nM CLM, 4 h
B
Median TPM
300
250
200
150
100
50
0
BRCA
Normal
SRSF1 SRSF2 SRSF3 SRSF7 PTBP1
COAD
Normal
LUAD
Normal
LUSC
Normal

Koo *et al.* Sup. Fig. 6

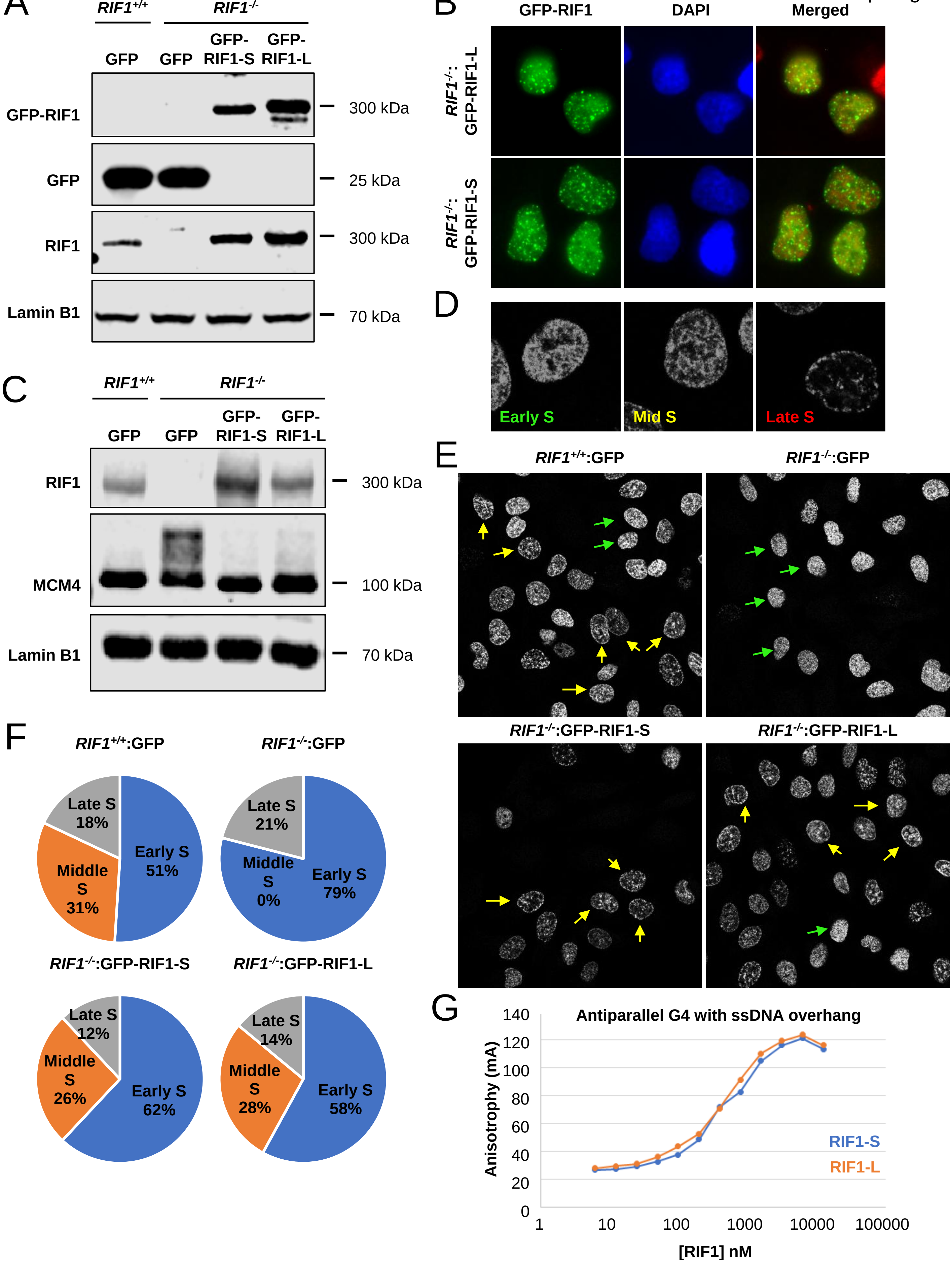

### Supplemental Videos

https://uwmadison.box.com/s/9rl8s2uyo2xu4pqjxvz0gq1iyss8tgoi

### Supplemental Tables

https://uwmadison.box.com/s/tcltyjbackeu0zotxs5h2mhpgc557v18
